# Plant functional groups and root traits are linked to exudation rates of mature temperate trees

**DOI:** 10.1101/2024.08.01.606171

**Authors:** Young E. Oh, Meghan G. Midgley, M. Luke McCormack, Katilyn V. Beidler, Marshall McCall, Savannah Henderson, Renato K. Braghiere, Richard P. Phillips

## Abstract

While root exudation has the potential to affect soil biogeochemistry profoundly, the process is rarely quantified in mature, field-grown trees. We measured rates of carbon (C) exudation in 11 trees species that exhibit divergent root traits, including gymnosperms and angiosperms that associate with either arbuscular mycorrhizal (AM) or ectomycorrhizal (EcM) fungi. Our goal was to explore how tree species, plant functional groups and root traits collectively influence exudation patterns. Intraspecific variation in exudation rates was larger than interspecific variation, and neither functional groups nor morphological traits alone could sufficiently explain variation in this flux. EcM-associated gymnosperms exuded 2.4 times more C than EcM angiosperms and 1.5 times more than AM gymnosperms. Exudation rates correlated positively with specific root length (SRL) and specific root area (SRA), and were correlated with root tissue density and root diameter in EcM-associated species. Mixed-effect models revealed that exudation rates were best determined by a combination of phylogenetic group, tree-mycorrhizal type and SRA, though a large portion of unexplained variation suggests that contemporary environmental and local edaphic conditions are likely important. Collectively, our results reveal that exudation is a complex physiological process governed by multiple factors and cannot be fully explained by functional groups or root traits alone. Instead, a combined consideration of these factors and new experimental approaches may be needed before exudation patterns can be linked to plant trait frameworks and incorporated into large-scale models.

## Introduction

Global environmental changes are altering plant community composition, with poorly understood impacts on belowground processes and biogeochemical cycling (Fei et al., 2017; Jo et al., 2019). Root carbon (C) exudation is a physiological process that links aboveground-belowground interactions (Bardgett, 2014; McCormack et al., 2017; Weemstra et al., 2022; Wen et al., 2022) and, in many cases, mediates ecosystem responses to global change (Norby et al., 2024; Phillips et al., 2011). Root exudates represent 5–20% of photosynthetically-fixed C (Chari et al., 2024), and much of this C fuels rhizosphere microbes that, in turn, determine soil organic matter dynamics (Chari & Taylor, 2022) and nutrient availability (Brzostek et al., 2014; Finzi et al., 2015; Meier et al., 2017; Yin et al., 2014). In this way, exudation rates affect ecosystem C balance through their effects on both nutrient uptake/primary production and microbial decomposition. Given this role, a deeper understanding of factors that mediate exudate fluxes should enhance our understanding of the ecosystem effects of plant community change (Freschet et al., 2021; Jo et al., 2019; McCormack et al., 2017).

Root traits impact many ecosystem processes (Bardgett, 2014), yet there is little consensus about which traits, if any, align with exudation rates. Exudation rates have been reported to be associated positively with both specific root length (SRL; Meier et al., 2020; Tückmantel et al., 2017; Wang et al., 2021) and negatively with root tissue density (RTD; Sun et al., 2017, 2021) - traits that capture different dimensions of root economic space (RES; Bergmann et al., 2020; McCormack & Iversen, 2019; Weemstra et al., 2016; Weigelt et al., 2021). In the RES, RTD and root N represent a ‘conservation’ gradient (e.g., ‘fast’ *vs.* ‘slow’), whereby long-lived, tissue-dense roots (high RTD) slowly provision N to hosts relative to fast-growing acquisitive roots with low RTD and high root N (Bergmann et al., 2020; Weigelt et al., 2021). Orthogonal to this axis is the ‘collaboration’ gradient (e.g., ‘outsourcing’ *vs.* ‘do-it-yourself’) defined by SRL and root diameter (Bergmann et al., 2020; McCormack & Iversen, 2019; Weemstra et al., 2016; Weigelt et al., 2021; Wen et al., 2022; Yaffar et al., 2022). Here, large diameter roots with low SRL are colonized by mycorrhizal fungi to a greater extent than thin, high SRL roots (Bergmann et al., 2020; Weigelt et al., 2021). However, links between exudation and both axes of the RES remain unclear, indicating that the relationship may depend on site factors (climate, soils and nutrient availability) and the traits of the species under consideration. As such, investigations of multiple tree species (with divergent traits) growing in a common soil may help resolve this apparent paradox.

Tree species also exhibit varying degrees of plasticity in terms of their root traits (Weemstra & Valverde-Barrantes, 2022), which could influence exudation dynamics. Traits like SRL and branching intensity (BI) are associated with nutrient acquisition (Comas & Eissenstat, 2009), and tend to be more plastic than traits linked to structural stability and longevity, such as RTD and root diameter <u>(Comas et al., 2012, 2014; Comas & Eissenstat, 2009; Sun et al., 2021)</u>. Whether exudation rates display greater intraspecific variation than morphological root traits remains unresolved <u>(Sun et al., 2021</u>), posing a key challenge for detecting the exudation-trait relationships.

Many root traits show a strong phylogenetic signal (<u>Brundrett, 2002; Comas et al., 2012, 2014)</u> suggesting that exudation rates may differ among tree species with divergent evolutionary histories (e.g., angiosperms *vs.* gymnosperms) and distinct mycorrhizal associations (e.g., arbuscular *vs.* ectomycorrhizal associations; AM *vs.* EcM). Moreover, if exudation patterns evolved as a means for dealing with nutrient limitations, links between tree species’ evolutionary history, root traits and exudation might be expected. Early gymnosperms had thick, dense, long-lived roots that associated with ‘ancestral’ AM fungi (Brundrett, 2002; Comas et al., 2012). As greater water and nutrient limitations emerged and selected for gymnosperms with highly-branched roots colonized by fungi derived from saprotrophs (i.e., EcM fungi), high exudation rates may have represented an additional strategy for nutrient acquisition (Brundrett, 2002; <u>Read & Perez-Moreno, 2003)</u>. When angiosperms arose in the early Cretaceous, some species evolved thin diameter, highly proliferative roots (Brundrett, 2002; Comas et al., 2012; Guo et al., 2008) whereas others - often in nutrient-poor soils - developed EcM associations (Comas et al., 2012; Read & Perez-Moreno, 2003). Whether exudation rates relate to tree species’ belowground C allocation and nutrient acquisition strategies is unknown, yet there are reasons to suspect that the evolutionary processes that shape root trait syndromes and tree-mycorrhizal associations also affect exudation.

There has been little consensus over whether exudation rates differ among tree species from different functional groups (Brzostek et al., 2013; Liese et al., 2018; Wang et al., 2021). Exudation could be greater in EcM trees (relative to AM trees) if exudation is a reflection of the C sink strength of roots, which is typically greater in EcM root systems (Hobbie, 2006). Alternatively, if exudation rates reflect C allocation tradeoffs within the root system (e.g., C exuded by roots comes at a cost to C used to support mycorrhizal fungi; Wen et al., 2019), one might expect higher exudation in AM trees where the C costs of supporting AM hyphae are low relative to EcM mycelium (Hawkins et al., 2023). To date, support for both hypotheses is apparent. In temperate forests, EcM trees have been shown to exude more C than AM trees (Brzostek et al., 2013; Phillips & Fahey, 2006; Yin et al., 2014), though this effect is not apparent in young trees (Liese et al., 2018) and the opposite pattern has been reported in sub-tropical forests <u>(Sun et al., 2021</u>). Likewise, deciduous trees have been shown to have higher exudation rates than evergreen trees in some temperate forests (Sun, et al., 2017; Wang et al., 2021) but not others (Brzostek et al. 2013). In a recent synthesis of dozens of studies, Chari et al. (2024) found no evidence of exudation differences between angiosperms and gymnosperms or between AM and EcM trees. These mixed findings highlight the need to investigate tree functional group effects on exudation at a common site where other factors (climate, tree age, soil characteristics, etc.) can be controlled for.

In this study, we assessed the effects of tree species, functional groups, and root traits on exudation rates in mature trees grown in monodominant plots in a common soil. Importantly, to disentangle the effects of functional groups (e.g., AM and EcM *vs.* angiosperms and gymnosperms) from root traits across the RES, we selected tree species from each functional group that spanned a range of root trait space. Our objectives were to (1) characterize the extent to which exudation rates vary among tree species and across functional groups, (2) determine which root traits in the RES, if any, are closely related to exudation rates, and (3) build a framework for predicting exudation using readily-measurable root traits and tree functional groups. We hypothesized that exudation rates would differ among tree species and functional groups due to differences in root traits, leading to the prediction that considering both root traits and functional groups would better predict exudation rates (H1). Additionally, we hypothesized that exudation is linked to one or more axes of the RES (H2): (a) congruent to the collaboration gradient (leading to the prediction that exudation correlates positively with SRL or SRA) or (b) congruent to the conservation gradient (leading to the prediction that exudation correlates negatively to RTD).

## Materials and methods

### Site description

This study was conducted in monoculture plots at the Morton Arboretum, Lisle, Illinois (41.81N, 88.05W). The plots were established between 1922 and 1948 to test and study “all the timber trees of the world which might come under consideration for reforestation purposes in this part of the country” (Morton Arboretum Staff, 1929). Soils in the plots are poorly drained Alfisols that form from a thin layer of loess (0.31 m) underlain by glacial till and Mollisols that formed from alluvium (Soil Survey Staff, NRCS, USDA, 2024). The soil series in the plots are primarily Ozaukee silt loams and Sawmill silty clay loam (Midgley & Sims, 2020). The area has a continental climate with temperatures ranging from -6°C in January to 22℃ in July and 800-1,000 mm mean annual precipitation.

Eleven tree species were selected to capture the heterogeneity in root traits among species from distinct functional groups: phylogenetic group (angiosperm *vs.* gymnosperm), tree-mycorrhizal association (AM *vs.* EcM), and leaf habit (deciduous *vs.* evergreen). Within each group, species were chosen based on mean SRL and root tissue N concentration (root N) - the traits that were found to correlate positively with exudation rates in previous studies (Meier et al., 2020; Sun et al., 2021; Wang et al., 2021). As such, selected eleven species spanned a wide range of SRL and root N for each group, ensuring that all selected species captured diverse trait space (Table 1). This allowed for minimizing phylogenetic covariations among traits while maximizing species trait dissimilarities. Out of the eight combinations, only two combinations were absent: evergreen-AM-angiosperms and evergreen-EcM-angiosperms (Table 1).

**Table 1.** List of eleven tree species in monodominant forestry plots at The Morton Arboretum (Lisle, IL, USA); All species are ≥70 years old except *Asimina triloba*.

| Species | Soil series | Soil order | Mycorrhizal type | Phylogenetic group | Leaf habit | Clade |
| --- | --- | --- | --- | --- | --- | --- |
| *Acer saccharum* | Ozaukee | Alfisol | AM | Angiosperm | Deciduous | Rosids |
| *Asimina triloba* | Ozaukee | Alfisol | AM | Angiosperm | Deciduous | Magnoliids |
| *Platanus occidentalis* | Ashkum | Mollisol | AM | Angiosperm | Deciduous | Eudicots |
| *Chamaecyparis pisifera* | Ozaukee | Alfisol | AM | Gymnosperm | Evergreen | Gymnosperms |
| *Juniperus chinensis* | Ozaukee | Alfisol | AM | Gymnosperm | Evergreen | Gymnosperms |
| *Taxodium distichum* | Sawmill | Mollisol | AM | Gymnosperm | Deciduous | Gymnosperms |
| *Carya ovata* | Ozaukee | Alfisol | EcM | Angiosperm | Deciduous | Rosids |
| *Quercus bicolor* | Sawmill | Mollisol | EcM | Angiosperm | Deciduous | Rosids |
| *Larix gmelinii* | Ozaukee | Alfisol | EcM | Gymnosperm | Deciduous | Gymnosperms |
| *Picea abies* | Ozaukee | Alfisol | EcM | Gymnosperm | Evergreen | Gymnosperms |
| *Tsuga canadensis* | Ozaukee | Alfisol | EcM | Gymnosperm | Evergreen | Gymnosperms |

*Note: AM, arbuscular mycorrhizal type; EcM, ectomycorrhizal type*

### Root exudation rates

Fine-root exudates were collected during the growing season of 2022 (i.e., from May to July 2022) using an *in-situ* culture-based cuvette system (Phillips et al., 2008). To mitigate the impact of variable weather, sampling campaigns were conducted under sunny and clear conditions, to the extent possible, and each plot was visited twice: once in late May/early June to collect exudates from three individuals and once in late June/early July to collect exudates from 3-4 additional individuals. The terminal roots were excavated carefully from the mineral topsoil below the organic layer. The excavated root segments were examined to ensure that the fine-root system consisted of the first three branching orders with an intact absorptive function. Organic matter and soil particles adhering to the root system were removed with DDI water with extreme caution while keeping the roots moist with wet paper towels. In cases where the distal fine roots were damaged or broken off, samples were discarded, and a new sample was prepared. The intact root systems were placed in cuvettes (30mL syringe) filled with sterile, C-free glass beads (>1mm diameter). The root systems with glass beads were flushed three times with C-free nutrient solution (0.5 mM NH_4_NO_3_, 0.1 mM KH_2_PO_4_, 0.2 mM K_2_SO_4_, 0.15 mM MgSO_4_, 0.3 mM CaCl_2_) to ensure the root segments and glass beads were well-mixed and to remove any C adhering to the root surface. To ensure the same amount of solution was added to the cuvettes, we added 15mL of nutrient trap solution in the field using a bottle-top dispenser. The cuvette was covered in aluminum foil to allow the root system to equilibrate with the cuvette environment. The same procedure was applied to the control (i.e., no root) cuvette with the same glass beads and nutrient solution. The cuvettes were placed at the exact excavated area and covered with soils and organic matter and incubated for approximately 24 hrs.

After the one-day incubation period, the sampled roots with the cuvette were clipped with care and brought to the laboratory for analysis. Within one hour of clipping, each cuvette was flushed with 15mL of the working nutrient solution three times to remove accumulated exudates in the cuvette. All solutions were filtered immediately through a sterile 0.22 μm syringe filter (Millex-GV 0.22µm PVDF 33mm Gamma Sterilized 50/Pk, Millipore Co., Billerica, MA) and refrigerated at 4°C until analyses (<24 h). All samples were analyzed for non-particulate organic C on a TOC analyzer (Shimadzu Scientific Instruments, Columbia, MD) within a day of sample collection. The total mass-specific exudation rate was calculated with the total C captured from the trap solution minus the total C flushed from the root-free control cuvettes divided by the dry root biomass and day (mg C * g _root_^-1^ * day^-1^).

### Root morphological and chemical traits

Roots originally placed in the cuvette were carefully collected from the cuvette, washed, and stored at 4°C until processing. Fine-root morphology was analyzed for all the fine roots with a transparent flat-bed scanner and the WinRHIZO program (Regent Instruments, Quebec, QC, Canada). Scans were collected at a resolution of 600 dpi. All root samples were dried at 65°C for at least 48 h, and the dried root biomass was used for root trait calculations. Specific root length (SRL, in m g^-1^ : the length of the fine roots divided by the corresponding root dry weight), specific root area (SRA, in cm^2^ g^-1^ : the area of the fine roots divided by the corresponding root dry weight), root tissue density (RTD, in g cm^-3^ : root dry weight divided by root volume), root branching intensity (BI, in the number of tips per total fine-root length), and root diameter (diameter, in cm) were calculated from WinRHIZO. Root N concentration (per dry weight) was measured independently in the lab using an elemental combustion system (Costech Analytical Technologies).

### Statistical analyses

We used an analysis of variance (ANOVA), mixed linear models, and variance partitioning to characterize the extent to which root exudation rates vary among tree species and across functional groups. To test for differences in exudation rates among tree species, we conducted pairwise comparisons after an ANOVA using a Tukey’s Honest Significant Difference (HSD) test. To test for differences in exudation rates among tree functional groups, we built a mixed linear model with mycorrhizal type, phylogeny, and their interaction as fixed effects and species-plot as a random effect using restricted maximum likelihood (‘lme4::lmer’ via REML). To evaluate the significance of each nested group in the model after accounting for all other groups, Type III ANOVA with Satterthwaite’s Method using the ‘lme4::anova’ was performed to summarize the results of each model. To control the likelihood of false positives in all linear mixed effects models, adjusted p-values from BH Correction (Benjamini-Hochberg) test were performed using the p.adjust function. To quantify the contributions of inter- vs. intraspecific variation to exudation rates in mixed effects models, a variation partitioning analysis was performed using the ‘VEGAN::varpart’. To show co-variations among root traits, a pairwise trait relationships between exudation rates and root traits were also performed using Pearson’s correlations at the individual tree level using ‘corr.test’ function. Root traits and exudation rates were natural-log-transformed prior to analyses to meet model assumptions of residual normality and homogeneity of variance.

To assess how and the extent to which root exudation rates are associated with root trait coordination, we used principal components analysis (PCA) (Weigelt et al., 2023) and Redundancy Analysis (RDA). To examine how exudation rates aligns with major dimensions in the PCA, we created an ordination of RTD, root N, SRL, Diameter, SRA, and BI along with exudation rates using princomp () with standardized PCA. To examine the significance of linear relationships between exudation rates and the first four axes, we created a PCA without exudation rates and preformed Pearson’s product-moment correlation test between PCs and exudation using cor.test function. To select the best predicting root trait or subset of predictors, we built a PCA with four core variables (RTD, root N, SRL, and Diameter) and evaluated the relationship between root exudation rates as a trait and the traits that comprise the PCA. We used RDA models for PCs to explain exudation using ‘VEGAN::rda’ and selected the best predicting trait using ‘VEGAN::ordistep’ with both forward and backward stepwise model selection.

To identify the functional groups and root traits that collectively best predict exudation rates, we used a stepwise model selection approach using linear mixed-effects models by ‘lme4::lmer’ via REML. The fixed effects included six root traits (RTD, root N, SRL, Diameter, SRA, and BI) along with mycorrhizal type or phylogenetic group. Monodominant plot identity (i.e., species-plot) was treated as a random effect. Model selection was based on improvements in Akaike Information Criterion (AIC) and likelihood ratio tests comparing full and reduced models. Building on the best-performing model, we further tested interactions between traits and functional groups (e.g., Exudation ∼ Trait × Functional Group). We examined the explanatory power of each model by calculating marginal (R²m) and conditional (R²c) R-squared values, where R²m represents variance explained by fixed effects and R²c includes both fixed and random effects (Nakagawa & Schielzeth, 2013). Model assumptions for selected models were verified via checks for residual normality, homoscedasticity, and unbiasedness. All statistical analyses were performed using R v.3.5.3 (R Core Team, 2017).

## Results

### Hypothesis 1: Exudation differences among species and functional groups

#### Variation in exudation rates among tree species and functional groups

We found partial support for H1, as species explained 22% of the total variation in exudation rates in our model (Adj. R^2^ = 0.22; p = 0.01). While not all species differed in their exudation rates (ANOVA using Tukey’s HSD test), *Larix gmelinii* exhibited significantly higher rates of root exudation compared to *Chamaecyparis pisifera* (p=0.03) and *Carya ovata* (p<0.01) (Fig. 1a; Table 2). The mean exudation rate of *L. gmelinii* (3.82 mgC g_root_^-1^ * day^-1^) was more than twice that of the second-highest species, *Picea abies* (1.52 mgC g_root_^-1^ * day^-1^) (Fig. 1a: Table 2).

**Fig 1.**
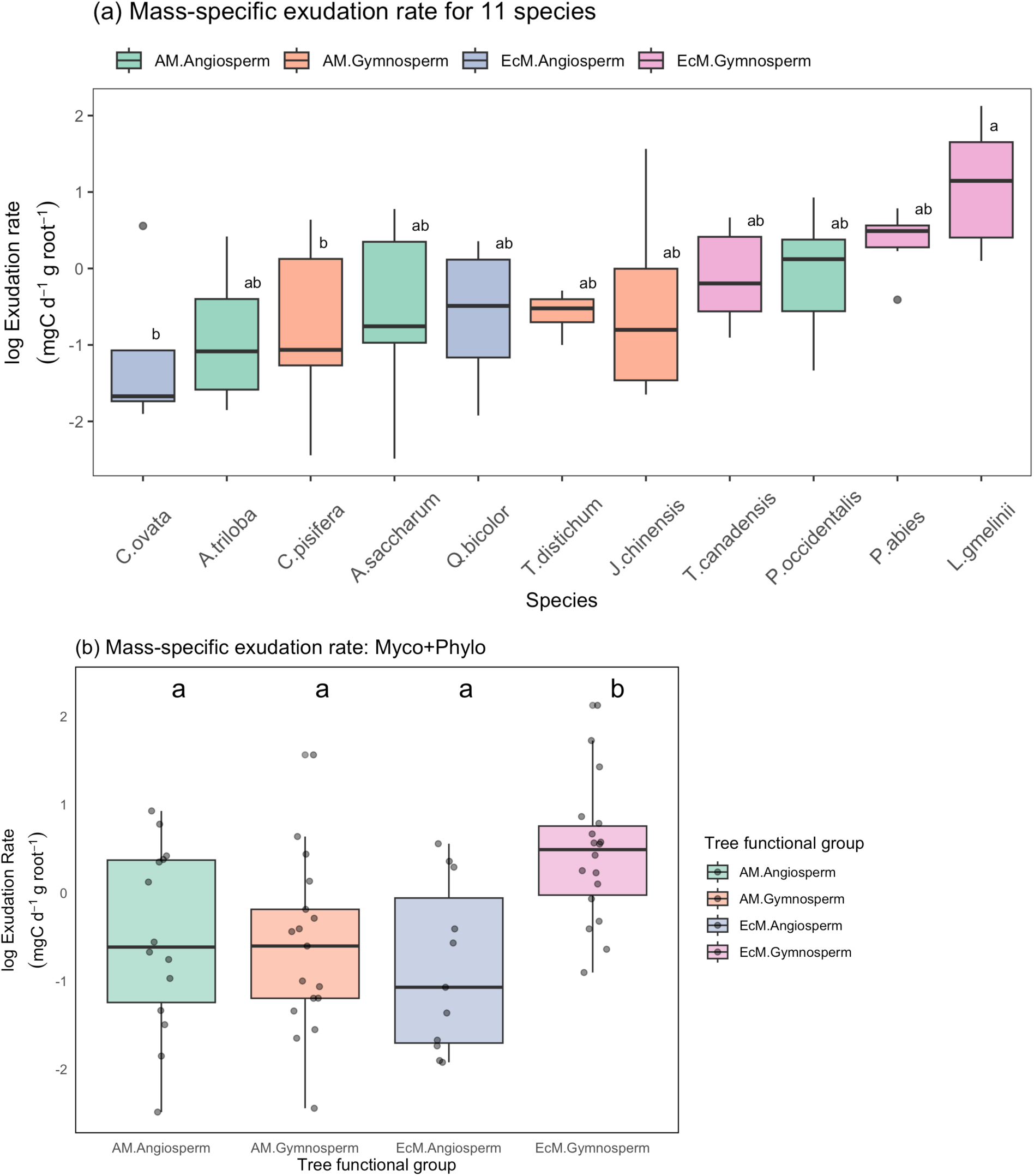
Boxplot of root exudation variation among: (a) 11 tree species and (b) combinations of mycorrhizal type (AM vs. EcM) and phylogenetic group (Angiosperm vs. gymnosperm) at The Morton Arboretum, Lisle, IL, USA. The central box in each boxplot represents the median and the interquartile range. The whiskers extend to the minimum and maximum value. In (a), different lowercase letters denote significant differences, as determined by a post hoc TukeyHSD test. In (b), different lowercase letters denote significant differences (Linear mixed model fit by REML; Std β = 9.6, p=0.03). Closed circles denote outliers.

**Table 2.** The species-specific means ± SDs of six core root traits at The Morton Arboretum, Lisle, IL, USA. n denotes the number of tree individuals sampled. Exudation: mass-specific exudation rate.

| Species (plot) | n | Exudation rate<br>(mg C g<sub>root</sub><sup>-1</sup> * day<sup>-1</sup>) | Specific root length<br>(SRL (cm g<sup>-1</sup>)) | Root diameter<br>(Diameter (mm)) | Root tissue density<br>(RTD (g cm<sup>-3</sup>)) | N concentration<br>(root N (%)) | Specific root area<br>(SRA (cm<sup>2</sup> g<sup>-1</sup>)) | Branching intensity<br>(BI (Tips cm<sup>-1</sup>)) |
| --- | --- | --- | --- | --- | --- | --- | --- | --- |
| *Acer saccharum* | 5 | 0.91 ± 0.87 | 1636.36 ± 529.41 | 0.34 ± 0.03 | 0.73 ± 0.14 | 1.23 ± 0.09 | 169.90 ± 42.69 | 3.19 ± 0.50 |
| *Asimina triloba* | 4 | 0.60 ± 0.63 | 1110.21 ± 104.32 | 0.64 ± 0.02 | 0.29 ± 0.05 | 2.91 ± 0.23 | 222.05 ± 28.12 | 0.72 ± 0.06 |
| *Platanus occidentalis* | 5 | 1.19 ± 0.88 | 1045.53 ± 540.38 | 0.52 ± 0.10 | 0.52 ± 0.07 | 1.74 ± 0.23 | 157.53 ± 52.86 | 2.95 ± 0.53 |
| *Chamaecyparis pisifera* | 7 | 0.75 ± 0.71 | 1528.93 ± 460.45 | 0.42 ± 0.06 | 0.51 ± 0.09 | 1.87 ± 0.18 | 195.13 ± 39.51 | 2.18 ± 0.56 |
| *Juniperus chinensis* | 6 | 1.22 ± 1.78 | 863.63 ± 273.33 | 0.55 ± 0.09 | 0.54 ± 0.11 | 1.39 ± 0.11 | 142.73 ± 25.23 | 2.40 ± 0.90 |
| *Taxodium distichum* | 4 | 0.58 ± 0.16 | 1696.22 ± 374.94 | 0.51 ± 0.08 | 0.31 ± 0.04 | 1.89 ± 0.26 | 264.06 ± 26.82 | 2.02 ± 0.42 |
| *Carya ovata* | 5 | 0.52 ± 0.69 | 1160.74 ± 209.51 | 0.37 ± 0.05 | 0.85 ± 0.17 | 1.25 ± 0.26 | 133.43 ± 25.23 | 5.32 ± 1.15 |
| *Quercus bicolor* | 6 | 0.73 ± 0.54 | 1439.00 ± 397.30 | 0.33 ± 0.02 | 0.86 ± 0.14 | 1.29 ± 0.14 | 146.67 ± 32.44 | 5.63 ± 0.48 |
| *Larix gmelinii* | 6 | 3.82 ± 2.83 | 975.76 ± 384.57 | 0.58 ± 0.11 | 0.43 ± 0.07 | 1.48 ± 0.12 | 167.97 ± 40.13 | 3.68 ± 0.60 |
| *Picea abies* | 6 | 1.52 ± 0.52 | 1639.29 ± 220.22 | 0.42 ± 0.04 | 0.45 ± 0.05 | 2.01 ± 0.13 | 214.35 ± 19.09 | 4.44 ± 0.66 |
| *Tsuga canadensis* | 6 | 1.05 ± 0.66 | 689.36 ± 276.65 | 0.59 ± 0.07 | 0.60 ± 0.17 | 1.24 ± 0.42 | 125.24 ± 46.82 | 4.16 ± 0.63 |

We also found a significant interaction between mycorrhizal type and phylogenetic group in exudation rates (Chisq = 9.27, Adj. p < 0.01; Table 3). On average, EcM gymnosperms exuded 2.4 times more C than EcM angiosperms and 1.5 times more C than AM gymnosperms (Fig. 1b). Despite these notable differences, tree mycorrhizal type, phylogenetic group, and leaf habit alone did not significantly explain variation in exudation (Chisq ≤ 0.98, Adj. p ≥ 0.48; Table 3). Additionally, we did not detect a significant interaction between mycorrhizal association and leaf habit (Chisq = 1.90, Adj. p =0.34; Table 3).

**Table 3.** Type III Analysis of Variance Table for predicting root traits of 11 tree species in The Morton Arboretum (n =60) using a linear mixed model (dependent variables∼myco x phylo x leaf + (1|species)). The species were nested in mycorrhizal types, phylogenetic groups and leaf habit. p-values (P<0.1) are highlighted in bold. The significance of fixed effects in the model was tested by Type III ANOVA with Wald chisquare tests. All measurements were log-transformed to ensure normality and improve homogeneity of variance.

| Fixed Effect | Mass-specific Exudation rate |  |  | Root diameter (mm) |  |  | Specific root length (m g<sup>-1</sup>) |  |  | Specific root area (cm<sup>2</sup> g<sup>-1</sup>) |  |  |
| --- | --- | --- | --- | --- | --- | --- | --- | --- | --- | --- | --- | --- |
|  | Chisq | Pr(>Chisq) | Adjusted p | Chisq | Pr(>Chisq) | Adjusted p | Chisq | Pr(>Chisq) | Adjusted p | Chisq | Pr(>Chisq) | Adjusted p |
| Mycorrhizal type (myco) | 0.98 | 0.32 | 0.48 | 4.37 | **0.04** | 0.11 | 0.08 | 0.77 | 0.77 | 2.44 | 0.12 | 0.16 |
| Phylogenetic group (phylo) | 0.02 | 0.88 | 0.92 | 0.08 | 0.78 | 0.78 | 1.15 | 0.28 | 0.34 | 3.92 | **0.048** | **0.10** |
| Leaf habit (leaf) | 0.01 | 0.92 | 0.92 | 0.09 | 0.77 | 0.78 | 1.45 | 0.23 | 0.34 | 4.83 | **0.028** | **0.08** |
| myco:phylo | 9.27 | **0.002** | **0.007** | 2.58 | 0.11 | 0.22 | 2.22 | 0.14 | 0.34 | 0.66 | 0.42 | 0.42 |
| myco:leaf | 1.90 | 0.169 | 0.337 | 0.09 | 0.77 | 0.78 | 1.27 | 0.26 | 0.34 | 2.26 | 0.13 | 0.16 |
| % of the total variance explained by random effect (Species plot) | > 0.1% |  |  | **55.2% (p<0.001)** |  |  | **33.3% (p<0.001)** |  |  | **26% (p<0.01)** |  |  |

| Fixed Effect | Root tissue density (g cm<sup>-3</sup>) |  |  | Root N concentration (%) |  |  | Branching intensity (cm<sup>-1</sup>) |  |  |
| --- | --- | --- | --- | --- | --- | --- | --- | --- | --- |
|  | Chisq | Pr(>Chisq) | Adjusted p | Chisq | Pr(>Chisq) | Adjusted p | Chisq | Pr(>Chisq) | Adjusted p |
| Mycorrhizal type (myco) | 9.42 | **0.0021** | **0.006** | 3.23 | **0.072** | 0.22 | 10.82 | **0.0010** | **0.0057** |
| Phylogenetic group (phylo) | 3.69 | **0.055** | **0.082** | 0.01 | 0.91 | 0.91 | 0.02 | 0.89 | 0.92 |
| Leaf habit (leaf) | 4.52 | **0.034** | **0.067** | 0.30 | 0.58 | 0.87 | 0.05 | 0.82 | 0.92 |
| myco:phylo | 0.42 | 0.52 | 0.52 | 0.13 | 0.72 | 0.87 | 0.59 | 0.442 | 0.88 |
| myco:leaf | 1.05 | 0.30 | 0.37 | 0.20 | 0.65 | 0.87 | 0.01 | 0.922 | 0.92 |
| % of the total variance explained by random effect (Species plot) | **45.5% (p<0.001)** |  |  | **55.4% (p<0.001)** |  |  | **74% (p<0.001)** |  |  |

Note. Numerator Degrees of Freedom = 1; Denominator Degrees of Freedom =60. Mass-specific exudation rate (mgC\*d<sup>-1</sup>\*g root<sup>-1</sup>)

Exudation rates exhibited over twice the variability of most root traits with a coefficient of variation (CV%) of 119% compared to lower variability across root traits (Table S1). This higher CV% corresponded to a high intraspecific variation (i.e., 77%) (Fig. S9; Table S3). Most root traits had CV%s below 40%, except for BI at 44% (Table S1), suggesting morphological traits are generally more conservative (i.e., less plastic) than exudation. High interspecific variation was observed in most root traits (RTD, root N, root diameter, and BI), whereas ‘composite’ and ‘acquisitive’ root traits such as SRL and SRA showed high intraspecific variability (>50%) and intermediate CV%s (38% and 30%, respectively) (Fig S9; Table S3). Together, unlike most root traits, exudation rates in our study can only be partially explained by species.

### Hypothesis 2: Exudation and the root economic space (RES)

The first two principal axes of the PCA generated with seven core root traits (RTD, root N, SRL, Diameter, SRA, BI, and exudation rates) explained 73% of the total variation in root traits (Fig. 2a; Table S5). The axis generated by RTD-root N was closely mapped onto PC1 reflecting the conservation gradient, while the axis generated by SRL-Diameter was loaded closely onto PC2 representing the collaboration gradient (Bergmann et al., 2020; Fig. 2a). The first five PCs demonstrated eigenvalues exceeding those predicted by random chance (as determined by Broken Stick analysis; Fig. S6), indicating that these axes accounted for more variance than would be expected under random conditions. In addition, the first three PCs showed eigenvalues greater than 1.0, indicating the significant contribution of three PCs to the ordination (Table S5; Tabachnik & Fidell, 1996).

**Fig 2.**
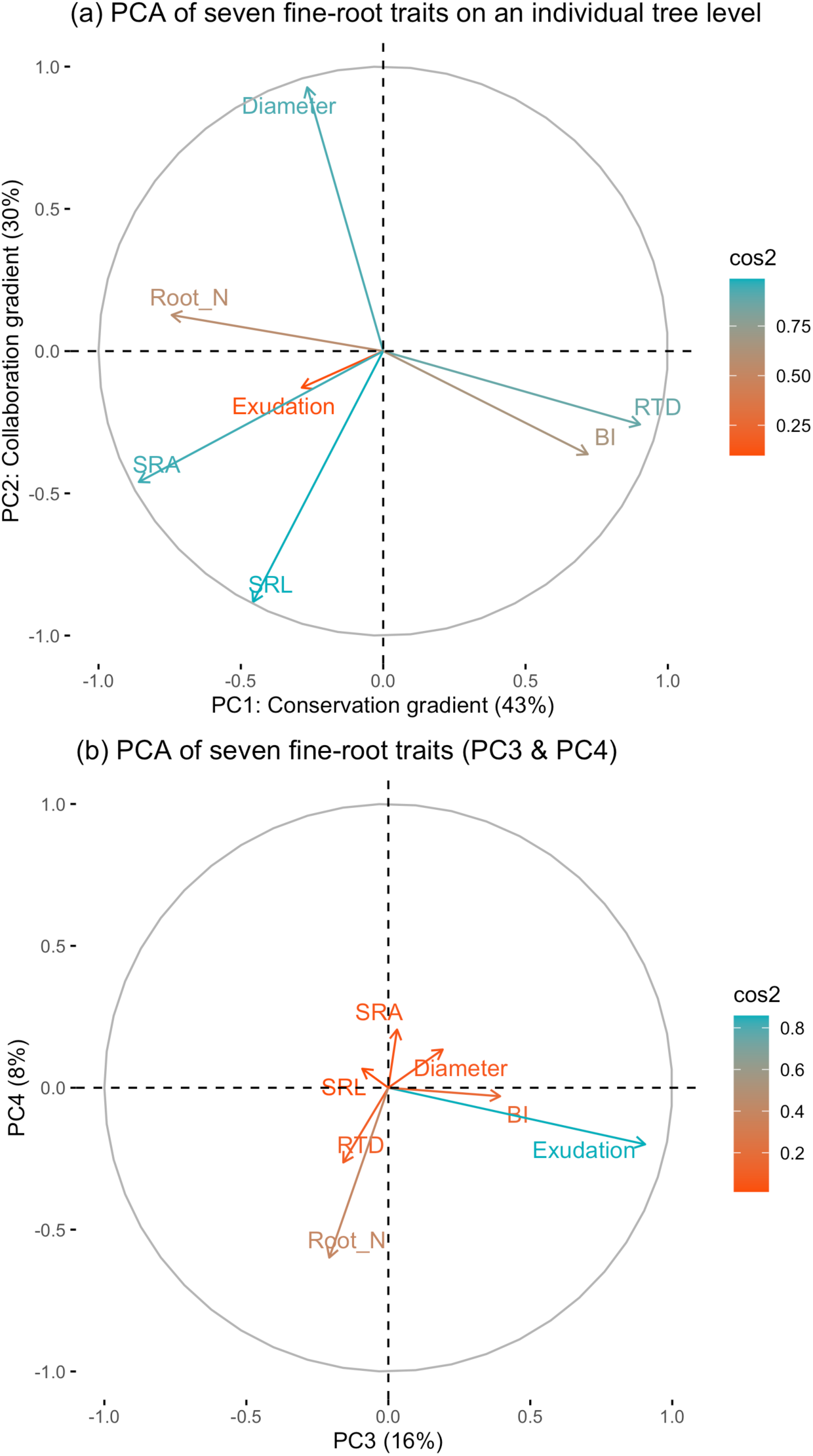
Standardized principal components analysis (PCA) of six fine-root functional traits and exudation rates on an individual tree level. ‘cos2’ values indicate that the quality of the representation of the variable on that principal component as a cos2 value close to 1 indicates that the variable is very well represented by the PC(s) in question. (a) PCA generated by six core root functional traits. PC1 and PC2 likely correspond to conservation and collaboration gradient respectively (Bergmann et al., 2020). (b) Two additional axes (PC 3 & 4) from the PCA.

Redundancy analysis (RDA) with permutation tests indicated that exudation rates were not significantly associated with variation along PC1 and PC2, which were generated with six root traits excluding exudation rates (p>0.15; Fig. S7). However, exudation rates were significantly correlated with PC3 (R² = 0.08, p=0.03) and PC4 (R² = 0.10, p=0.02), while the two PCs accounted for 10% and 6% of the total variation, respectively (Fig. S7). PC3 and PC4 together explained a greater proportion of variance in exudation (exudation ∼ PC3 + PC4; R² = 0.18, p=0.004). Notably, stepwise model selection based on RDA of the PCA derived from four core root traits (SRL– Diameter, RTD–Root N; Fig. S4) identified RTD as the only significant predictor of exudation (AIC = 3.56, F = 5.05, p=0.015). These results suggest that exudation likely correlates with the conservation gradient, but also suggest that root exudation is more strongly associated with trait variation captured by more than just the first two principal components in the RES.

### The effects of tree functional groups on trait-exudation relationships

#### Mixed-effect model predictions

Consistent with the prediction derived from the first hypothesis, incorporating both root traits and tree functional groups enhanced model predictions of exudation. As such, the best-performing mixed-effects model included SRA and the interaction term between the phylogenetic group and mycorrhizal types (exudation∼ SRA + mycorrhizal-type:phylogeny + (1|species); R^2^c = 0.36; p<0.01; Fig. 3; Table 4). That is, tree species with higher SRA - indicative of more acquisitive root strategies - exhibited significantly higher exudation rates (Std β = 1.13 ± 0.39, Adj. p=0.013; Fig. 3; Table 4). However, the interaction between mycorrhizal-type and phylogeny modified this relationship: compared to the baseline group (EcM-Gymnosperms), all other combinations (AM-Angiosperms, EcM-Angiosperms, and AM-Gymnosperms) showed significantly lower exudation rates (Adj. p=0.01; Fig. 1b). A larger model that additionally included root N also significantly predicted exudation rates (p<0.01; Table S4), albeit with a slightly reduced fit. These results supported our second hypothesis, suggesting that exudation rates might be influenced by both acquisitive and conservative root traits, as captured by variation in SRA and root N.

**Fig 3.**
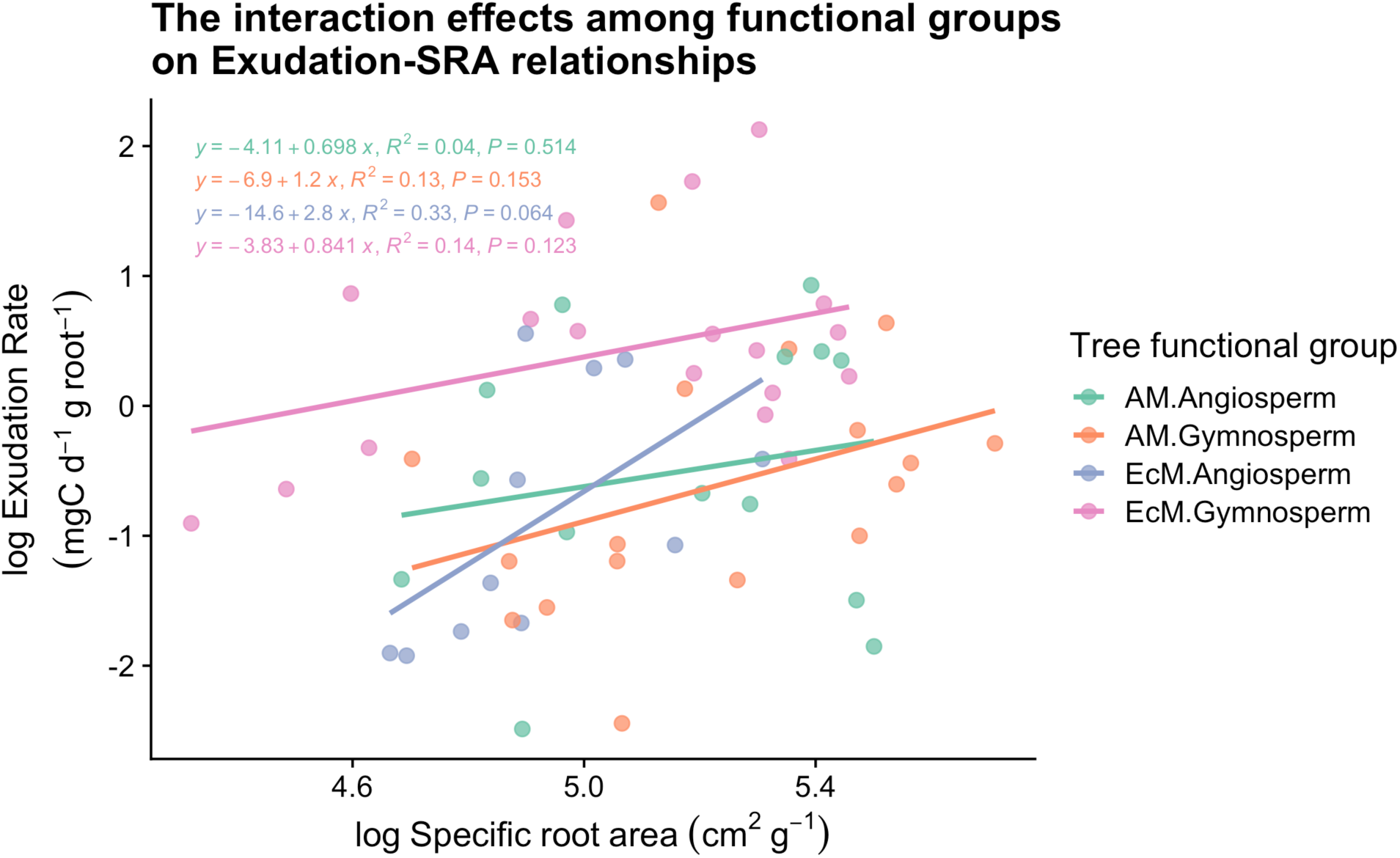
Interaction effects of tree functional groups on the relationship between log-transformed specific root area (SRA) and mass-specific exudation rates. The model depicts the interaction effects described in the best-predicting model selected by a stepwise model selection approach: Exudation ∼ SRA + myco.type:phylogeny + (1|species) (Table 4). Linear regression lines are shown for each group combination defined by mycorrhizal association (AM or EcM) and phylogeny (Angiosperm or Gymnosperm). Each equation displays the linear fit, R², and p-value. Among groups, EcM Angiosperms showed the steepest positive relationship between SRA and exudation rates (R² = 0.33, P = 0.064), while other groups exhibited weaker or non-significant trends. This interaction reflects differential trait-based exudation patterns across mycorrhizal and phylogenetic strategies.

**Table 4.** Mixed effects models for the effects of mycorrhizal type on exudation-to-trait relationships. The model considered root traits × mycorrhizal type + (1|species.plot) to predict exudation rates. ‘species.plot’ represents species-specific monodominant plots at The Morton Arboretum, Lisle, IL. The best model (exudation ∼ SRA + Mycorrhizal type:Phylogeny) is selected via a stepwise reduction approach using mixed models. The other models are ordered from the lowest AIC values. The numbers in the table represent coefficients estimate (Std β) and standard error (Std SE) in brackets. Adjusted p-values were calculated using the Benjamini-Hochberg (BH) correction for multiple testing. Bold value indicates statistical significance of Adjusted p-values less than 0.05. Significance levels of Adjusted p-values: *p<0.05; **p<0.01; ***p<0.001. The comprehensive table for the effects of phylogeny on trait-exudation relationships is in Table S2.

| Model to predict exudation rates<br>- AIC values | Significant Fixed effect term<br>- $R^2_m$ & $R^2_c$ | Std $\beta$ (Std SE) | Adjusted p |
| --- | --- | --- | --- |
| SRA + Mycorrhizal type : Phylogeny | (Intercept) | -5.2936 (1.99)* | **0.013** |
| Best model selected with AIC =164.8 | SRA | 1.13 (0.39)** | **0.013** |
|  | AM : Angiosperm | -1.05 (0.33)* | **0.013** |
|  | EcM : Angiosperm | -1.13 (0.35)* | **0.013** |
|  | AM : Gymnosperm | -1.23 (0.33)** | **0.013** |
| | - $R^2_m = 0.34$; $R^2_c = 0.36$ | | |
| RTD * Mycorrhizal type | RTD : EcM | -2.05 (0.64)** | **0.008** |
| - AIC=163.2 | - $R^2_m = 0.24$; $R^2_c = 0.44$ | | |
| SRA * Mycorrhizal type | SRA | 1.76 (0.64)* | **0.023** |
| - AIC=169.3 | - $R^2_m = 0.33$; $R^2_c = 0.33$ | | |
| SRL * Mycorrhizal type | SRL | 1.54 (0.45)** | **0.005** |
| - AIC=170.7 | - $R^2_m = 0.19$; $R^2_c = 0.43$ | | |
| Diameter (mm) * Mycorrhizal type | ECM | 2.91 (0.90)* | **0.016** |
| - AIC=173.0 | Diameter : ECM | 3.09 (1.09)* | **0.027** |
| | - $R^2_m = 0.10$; $R^2_c = 0.27$ | | |

Note) Insignificant models (i.e., Root N \* Mycorrhizal type; BI \* Mycorrhizal type) and fixed effect terms are not reported in this table. See Table S2 for comprehensive results that include the effects from phylogeny on trait-exudation relationships. All measured variables were log-transformed to ensure normality of data. (Intercept) represents baseline exudation rate (AM reference). $R^2_m$ , Marginal $R^2$ ; $R^2_c$ , Conditional $R^2$ , RTD. $R^2_m$ represents variance explained by fixed effects only, whereas $R^2_c$ indicates variance explained by both fixed and random effects. Root Tissue Density ( $\text{g}/\text{cm}^3$ ); SRA, Specific Root Area ( $\text{m}^2/\text{g}$ ); Specific Root Length ( $\text{cm}/\text{g}$ ); Diameter, root mean diameter (mm); Root N, Root N Concentration (%); BI, Branching Intensity ( $\text{cm}^{-1}$ ).

#### Exudation-trait correlations in mixed effects models

Linear mixed effects models that included significant effects of mycorrhizal association on exudation provide partial support for exudation as both an acquisitive resource exploitation strategy and a physiological process governed by C allocation tradeoffs. Across all species, variation in SRL, and SRA, and partly in root diameter, significantly accounted for variation in root exudation (Fig. 4; Table 4). The negative relationship of exudation with RTD and positive relationship with root diameter were modulated by mycorrhizal association (Fig. 4; Table 4), while phylogenetic group showed little effects on trait-exudation relationships (Table S2).

**Fig 4.**
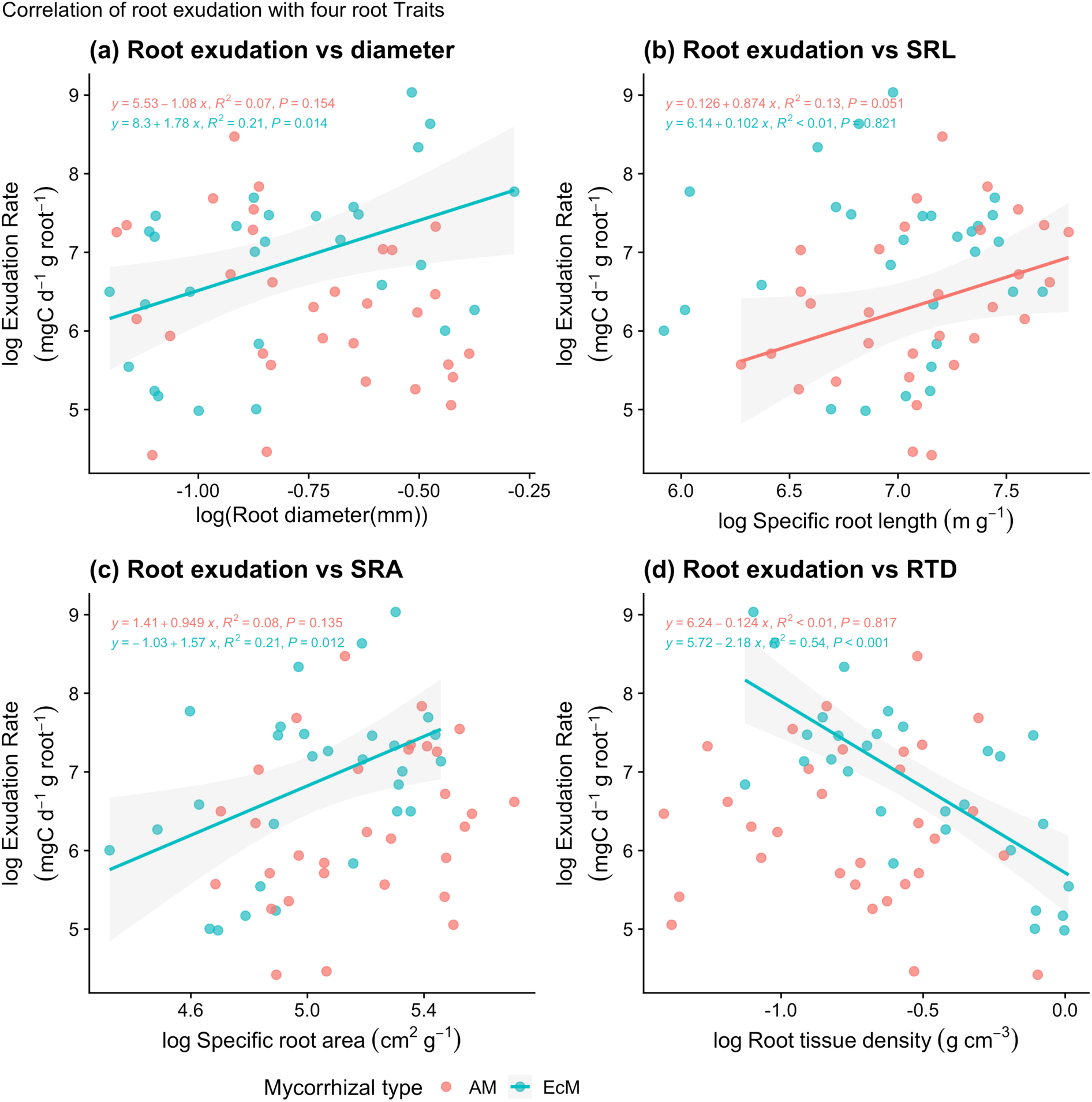
Linear relationships of log-transformed root exudation with (a) root diameter, (b) SRL, (c) SRA, and (d) RTD measured at The Morton Arboretum, Lisle, IL, USA. The fitted lines indicate statistically significant relationships of root exudation with root traits (p<0.1) that are nested with AM and EcM mycorrhizal association.

Specifically, high SRL and SRA were positively correlated with exudation rates across species (SRL: Std β = 1.5, Adj. p<0.001; SRA: Std β = 1.8, Adj. p=0.03; Table 4). Unlike SRA, this positive relationship between exudation and SRL appeared stronger among AM trees, as indicated by a marginally significant linear relationship in a simple linear model nested in mycorrhizal association (R² = 0.13, p=0.051; Fig 4b). The marginally significant interaction term (SRL × Mycorrhizal Type, Std β = -1.09, Adj. p=0.11; Table 4) suggests that EcM association may modulate this relationship, potentially exhibiting a weaker or even negative association between SRL and exudation compared to AM trees. However, further study is needed to confirm these effects, especially since the relationship was not detected in the bi-variate analysis (Pearson’s correlation ‘r’; Fig S1).

While no trend between exudation and RTD across species was detectable, RTD and exudation showed a significant interaction (Std β = -2.0, Adj. p=0.001, Table 4), showing that in ECM trees, increasing RTD was associated with a large decrease in exudation rates (Fig. 4d), with this relationship being more pronounced in gymnosperms (Table 3; Fig 1b). Exudation generally decreased with root diameter across species with marginal significance (Std β = -1.5, Adj. p=0.06; Fig. 4a; Table 4). Notably, EcM trees exhibited higher exudation rates than AM trees after controlling for diameter (Std β = 2.9, adjusted p=0.01; Fig 4a; Table 4), leading to a significant interaction between tree mycorrhizal association and diameter. Together, exploitative root traits such as SRA and SRL associated positively with exudation across species while EcM association significantly influenced the relationships of exudation with RTD, SRA and root diameter.

## Discussion

Our study aimed to identify key drivers of mass-specific root exudation in mature field-grown trees. We hypothesized that exudation rates would differ among tree species and functional groups (H1). Further, we hypothesized that exudation rates would be associated with at least one of the axes of the RES (H2): along the collaboration gradient of the RES owing to exudation’s functional role as a nutrient acquisition strategy and/or along the conservation gradient of the RES owing to exudation’s role as a competing sink for C (e.g., root tissue construction costs). We found partial support for H1, as exudation rates varied partly among tree species, and EcM gymnosperms exhibited greater exudation rates than other tree functional groups. In partial support of our second hypothesis, exudation rates correlated positively with SRL and SRA (across all species) and in EcM trees, correlated with conserved traits such as root diameter and RTD. However, while exudation rates were loaded weakly onto the ‘fast’ side of the conservative gradient in the RES, exudation was not correlated with the first two axes. Rather, exudation was better predicted by independent third and fourth (i.e., non-RES) axes. Finally, we found that the best model to predict exudation rates across all 11 species contained SRA (with a significant correlation with SRL) coupled with a strong influence from mycorrhizal type on phylogeny. Collectively, our study indicates that root exudation is a complex physiological process and cannot be fully explained by species identity or root traits alone. Instead, a combined consideration of these factors offers a more accurate prediction of fine-root physiological functioning.

### Tree functional groups partially account for variation in exudation

EcM gymnosperms generally had the highest exudation rates (two-fold higher than other groups), though the reasons remain elusive. High exudation rates have been reported for other EcM gymnosperms in temperate forests (Abramoff & Finzi, 2016; Brunn et al., 2022; Jiang et al., 2021; Yang et al., 2020; Zhang et al., 2016) and under controlled conditions (Sell et al., 2022; Yin et al., 2013). One of the highest exuders in our study, *Larix gmelinii*, has been reported to have elevated exudation rates in a boreal forest as well (Yin et al., 2023). Although the influence of EcM associations on root traits is well documented, their differential impacts on root traits and exudation between angiosperms and gymnosperms remain unclear.

Three potential mechanisms driving this pattern include tradeoffs between C allocation to EcM fungi and root exudates, leakiness of highly colonized gymnosperm roots, and high exudation rates of EcM fungi themselves. First, EcM gymnosperms might allocate more C to the rhizosphere and lesser to their mycorrhizal partners. For instance, a global synthesis by Hawkins et al. (2023) reported that C allocation to EcM mycelium biomass was ∼2.5 times greater in broad-leaved trees (mostly angiosperms) than needle-leaf trees (mostly gymnosperms). However, the <u>Hawkins et al</u>. <u>(2023)</u> synthesis did not account for biome differences, suggesting that reports of lesser C allocation to mycelium in EcM gymnosperms may have been confounded by climate (e.g., if boreal forest needle-leaf trees were compared to temperate/tropical forest broad-leafed trees). In fact, some of the most striking examples of prodigious EcM mycelium come from studies of EcM gymnosperms like *Pinus spp.* (Anderson & Cairney, 2007), and EcM gymnosperms tend to possess short, thick roots (Fig S8) that are well-colonized by mycorrhizal fungi (Cheng et al., 2016; Comas & Eissenstat, 2009; Ma et al., 2018). As such, a second factor that could explain greater exudation in EcM gymnosperms is mycorrhizal-root leakiness. Little is known about whether the mycorrhizal roots of EcM gymnosperms differ from those of EcM angiosperms in leakiness (Farrar et al., 2003). If the magnitude of leakiness is determined by the strength of the C sink - as described by the “hole in the pipe model” for nitrous oxide gas in Firestone and Davidson (1989) - greater exudation in EcM gymnosperms would result from greater C allocation to mycelium. Finally, while EcM fungi are a sink for plant-derived C (Prescott et al., 2020), they too can exude organic C (e.g., oxalic acid), resulting in an additional source of C in the cuvettes (Ahonen-Jonnarth et al., 2000; Sun et al., 1999; Van Schöll et al., 2008). Further studies are required to elucidate the mechanisms that drive high exudation rates of EcM gymnosperms.

### The relationships between exudation and root traits that comprise the RES

Exudation rates were aligned weakly with the fast side of the ‘conservation’ gradient defined by RTD and root N in the RES while also showing a strong association with SRA (Sun et al., 2021). These results suggest that exudation may be linked to acquisitive root traits (e.g., high SRA and root N) to optimize soil resource uptake (Hypothesis 2a; Eissenstat, 1991; Eissenstat & Yanai, 1997; Lv et al., 2023; Weemstra et al., 2016). These patterns may also reflect a crucial root-soil process where rapid exudation increases soil N availability (Meier et al., 2017), thereby enhancing root metabolic activity characterized by low RTD and high root N (Sun et al., 2021; Wen et al., 2022). We also found that RTD was the single significant predictor for exudation in the RES, partially supporting the prediction that exudation could correlate with root traits related to the conservation gradient reflecting structural stability, longevity, and metabolic activity such as RTD and root N (Hypothesis 2b). The observed significant relationships of exudation with RTD (from RDA analysis) and SRA (from mixed-effects models) likely result from the interplay of root area, tissue permeability, and diffusion strength - all of which can influence exudation rates (<u>Akatsuki</u> <u>& Makita, 2020;</u> Farrar et al., 2003; Jones et al., 2009; <u>Lv et al., 2023; Sun et al., 2021</u>). Since higher RTD often is association with reduced root leakiness and metabolic activity (Farrar et al., 2003), roots with less dense tissues, high root N, and larger surface areas are more likely to exude C. Consistent with our findings, recent studies in cool-temperate and subtropical forests have also reported significant relationships of exudation rates with RTD (-), root N (+), and SRA (+), but not SRL (Akatsuki & Makita, 2020; Sun et al., 2021).

The alignment of exudation within the RES suggests that exudation may function as an alternative soil resource exploitation strategy (Sun et al., 2021; Wen et al., 2022), modulated by the interplay between mycorrhizal associations (Brzostek et al., 2013) and root traits under phylogenetic constraints (Williams et al., 2022). However, exudation rates were not associated with the first two PC axes, and additional PC axes (PC3 and PC4; Fig. 2b) were needed to explain variation in exudation. This suggests that exudation may be more strongly associated with traits not considered or other factors. The RES simplifies trait space by collapsing trait variation into ‘collaboration’ and ‘conservation’ gradients (Bergmann et al., 2020), yet other dimensions of plant trait variation (e.g., plant height and rooting depth) were found to exist (Weigelt et al., 2021). Whether exudation aligns with this third axis of plant variation is unknown (Freschet et al., 2021; McCormack et al., 2017; Weemstra et al., 2022) or links to other unmeasured processes (e.g., C assimilation and allocation) is not well-understood, but this would be a fruitful line of inquiry in future studies.

### Trait-to-exudation relationships constrained by functional group interactions

Exudation rates were significantly linked to acquisitive root traits such as high SRA, high SRL, low RTD, while their variability was strongly influenced by EcM association, root morphology, and C construction costs. Notably, the model that included the interaction between mycorrhizal association and phylogenetic lineage in predicting exudation–SRA relationships outperformed all others (Table 4). These findings collectively suggest that exudation may be linked to acquisitive strategies (e.g., high SRL and SRA, narrow diameters, and low RTD) that favor foraging efficiency under the control of both mycorrhizal association and evolutionary history (Hypothesis 2a; Eissenstat, 1991; Eissenstat & Yanai, 1997; Lv et al., 2023; Weemstra et al., 2016).

Interestingly, we observed that EcM association influenced the relationship between exudation and diameter, where exudation rates increased with root diameter. This trend was predominantly driven by EcM gymnosperms, which exhibited short, thick roots (Fig. S8) and high exudation rates (Fig. 1b) - highlighting their reliance on symbiotic partners for nutrient acquisition. However, a previous study measuring exudation rates of EcM gymnosperms (*Pinus* and *Larix spp.*) found a significant negative relationship between exudation rates and root diameter (Akatsuki & Makita, 2020). These divergent results suggest that while the associations with EcM do broadly affect plant exudation rates, the direction of those relationships likely depends on several factors, including species identity, trait variations and methodological differences (Williams et al., 2021). For example, Akatsuki & Makita (2020) used a glass-fiber filter method that targeted smaller root segments (< 3rd order), potentially <u>capturing higher mass-specific exudation rates than the cuvette-base</u> <u>methods of Phillips et al. (2008)</u>. This suggests that EcM gymnosperms exude root carbon mostly from lower-order roots and that our <u>cuvette-base methods</u> may underestimate mass-specific exudation rates owing to inclusion of larger root segments (Akatsuki & Makita, 2020). Together, this highlights the complexity of trait-to-exudation relationships and the need for further studies to uncover generalizable patterns across diverse ecosystems.

Unlike EcM trees, which exhibit differences in some root traits across phylogenies, AM association did not significantly predict most root traits and trait-exudation relationships in our study (Fig. S8) (Akatsuki & Makita, 2020). Instead, AM angiosperms demonstrated greater within-group variability, including exudation rates (Fig. S8), suggesting a flexible, ‘do-it-yourself’ resource acquisition strategy with high plasticity (Bergmann et al., 2020). Together, these findings indicate that elevated exudation rates can result from (1) the interplay of EcM-associated root traits and phylogenetic constraints or (2) enhanced intraspecific variability in acquisitive root traits, allowing for efficient soil resource exploitation with moderate exudation rates.

### Challenges and opportunities

Exudation rates for mature, field-grown trees are rare, as all methods used to measure exudation introduce some artifacts; our study is no exception. While the root cuvette method of Phillips et al. (2008) has been used in dozens of forest studies (Chari et al., 2024), important methodological questions remain about how to minimize damage to root systems before placement into cuvettes, how to normalize the rates measured in different scales, the chemistry and sterility of the trapping solution, the duration of the equilibrium period, the duration of the collection period, and the contribution of mycorrhizal fungi to the total exudates (Oburger & Jones, 2018). Moreover, the high degree of spatio-temporal variation in exudation (Jacoby et al., 2017; Jiang et al., 2021; Yang et al., 2020) affects how many samples are needed to draw inferences about species-specific patterns and trait-exudation correlations. While we attempted to minimize variability by selecting tree species grown in a common soil, applying the same methodology to all trees, etc., the substantial amount of variability in exudation within species suggests that this flux may be better linked to dynamic physiological processes such as C assimilation and allocation than to (relatively) static morphological traits.

## Conclusion

We revealed that root exudation was positively associated with acquisitive root traits (SRL and SRA), while exudation-trait relationships were modulated by mycorrhizal association and phylogenetic lineage. EcM trees appeared to influence trait–exudation relationships, especially for RTD and root diameter, with these effects being more pronounced in gymnosperms. As such, constraining trait-exudation relationships with tree functional groups did improve model predictability. Importantly, we demonstrate that root exudation may be a complex physiological process that cannot be explained by species identity, functional groups or individual root traits alone. High intraspecific variation in root exudation (unlike stable morphological traits) likely contributed to the weaker alignment of exudation with the RES. Instead, exudation was linked to additional functional axes beyond the first two axes in the RES. Given the emerging interest in including root physiological traits into the RES (which is based mostly on morphological and chemical traits), our findings suggest that incorporating such dynamic processes into the RES may pose significant challenges and require to identify additional drivers of such dynamics other than root traits. As more mechanistic studies are developed (e.g. tracking exudate responses to tree girdling or nutrient solution culture alterations) and methods for capturing exudates are improved, our ability to understand the role and function of exudates in the environment should come into greater focus.

## Acknowledgements

This study was supported by the Center for Tree Science fellowship from The Morton Arboretum and Indiana University-Bloomington. Funding to YO and RPP was provided by NSF, DEB, MacroSysBIO & NEON-Enabled Science (Award# 2106096). We thank Elizabeth Huenupi for facilitating lab work in Indiana University and The Morton Arboretum Soils Lab REU students for lab assistance. We are grateful for constructive comments provided by Phillips Lab members.

## Author contributions

M.G.M., K.V.B., M.L.M., Y.E.O. and R.P.P. designed the study; M.G.M., K.V.B., M.L.M., Y.E.O., M.M., R.K.B., and S.H. performed the research; M.G.M., M.M., and Y.E.O. analyzed the data; Y.E.O., M.G.M. and R.P.P. wrote the paper with input from all authors.

