## Supporting_Information for "Plant functional groups and root traits are linked to exudation rates of mature temperate trees"

The following Supporting Information is available for this article:

**Fig. S1)** Pearson's correlation coefficient matrix chart showing trait-covariation

**Fig. S2)** The phylogenetic tree of 11 tree species at The Morton Arboretum, Lisle, IL, USA.

**Fig. S3)** The relationships of the total exudation rate with (a) total root mass (g) and (b) total root surface area ($cm^2$)

**Fig. S4)** (a) Root economic space (RES) projected by four core fine-root functional traits. (b) RES on an individual level with seven root traits and the 95% confidence ellipses of four nested combinations of tree functional groups (mycorrhizal type + phylogenetic group)

**Fig. S5)** Phylogenetically informed principal component analyses (PCA) of seven core traits of 11 mature tree species (n=11), (a) the broken stick model test results, the relationship of exudation rates with (b) the conservation gradient (PC1), (c) the collaboration gradient (PC2), and PC3 using linear model

**Fig. S6)** Statistical test results of principal components of the RES projected by standardized PCA of six root traits and exudation rates (Fig 4)

**Fig. S7)** Relationships between root exudation rates and (a) PC1 (conservation gradient), (b) PC2 (collaboration gradient), (c) PC3 (undefined), and (d) PC4 (undefined) of the RES based on six root traits excluding exudation rates.

**Fig. S8)** Variations of root traits ((a) Branching intensity, (b) RTD, (c) Root N, (d) SRL, (e) diameter, and (f) SRA)

**Fig. S9)** Variance partitioning of fine-root functional traits of 11 tree species.

**Table S1)** Descriptive statistics of the eight fine-root traits of eleven tree species (n=60)

**Table S2)** Comprehensive mixed effects models for the effects of mycorrhizal type on exudation-to-trait relationships

**Table S3)** Variance partitioning of eight fine-root functional traits

**Table S4)** Best predicting Mixed effects models of fine-root morphological traits on mass-specific exudation rate

**Table S5)** Results table of principal component analysis (PCA) on the six root traits, mass-specific exudation, and N uptake rates from 11 tree species

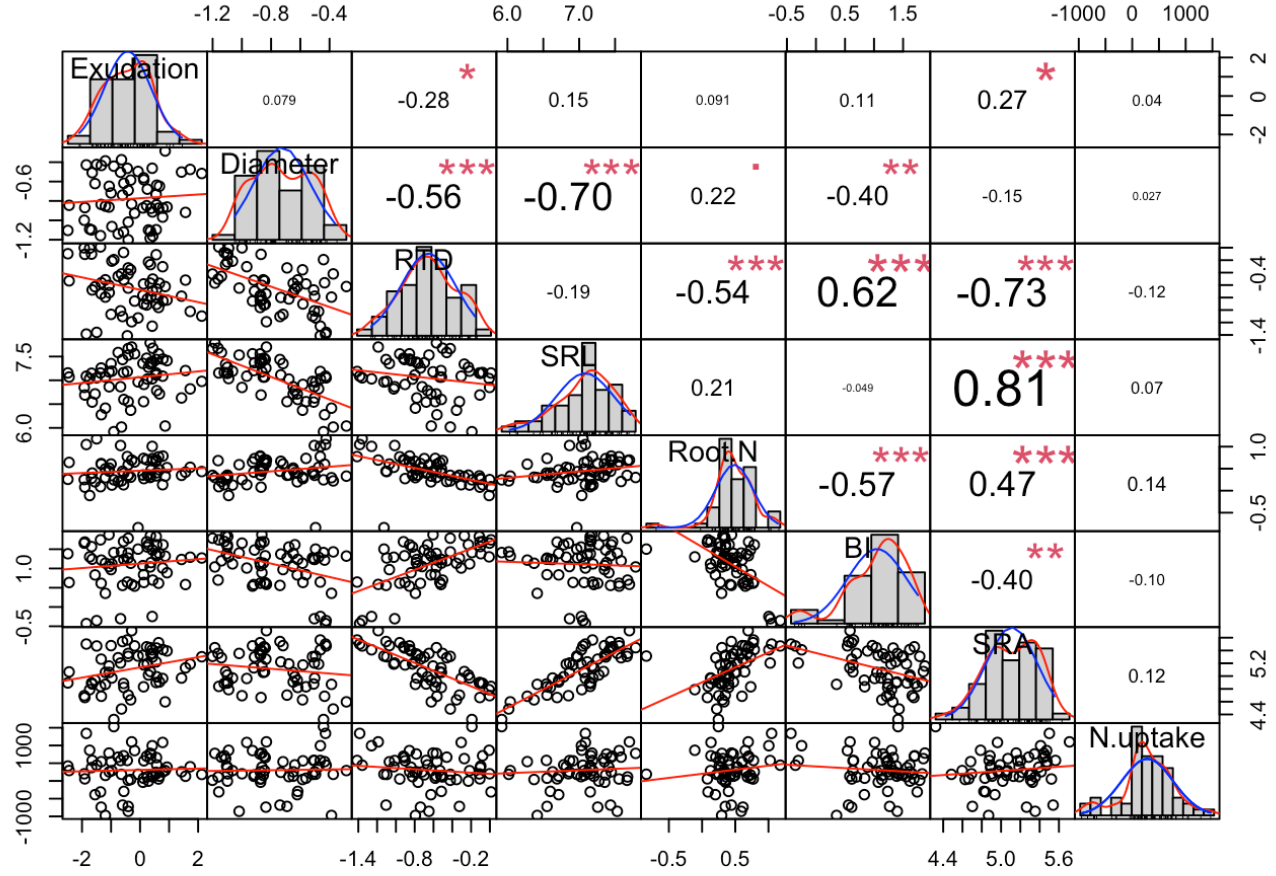


Fig S1) Pearson's correlation coefficient matrix chart with 95% confidence interval on individual tree level (n=60) among eight fine-root traits occurring 11 different monospecific mature-tree plots at The Morton Arboretum, Lisle, IL, USA. Significance level of correlations was denoted as ***, p<0.001;*** , *p<0.01;* , p<0.05. Exudation, mass-specific exudation rate ($m$gC d$^{-1}$ g$_{\cdot root}^{-1}$); Diameter; fine-root diameter (mm); RTD, root tissue density (mg cm$^{-3}$); SRL, specific root length (m g$^{-1}$); Root [N], root nitrogen concentration (%); BI, branching intensity (tips per cm); SRA, specific root area (cm$^{2}$ g$^{-1}$); N.uptake, mass-specific net inorganic N uptake ($m$gN d$^{-1}$ g$_{\cdot root}^{-1}$).

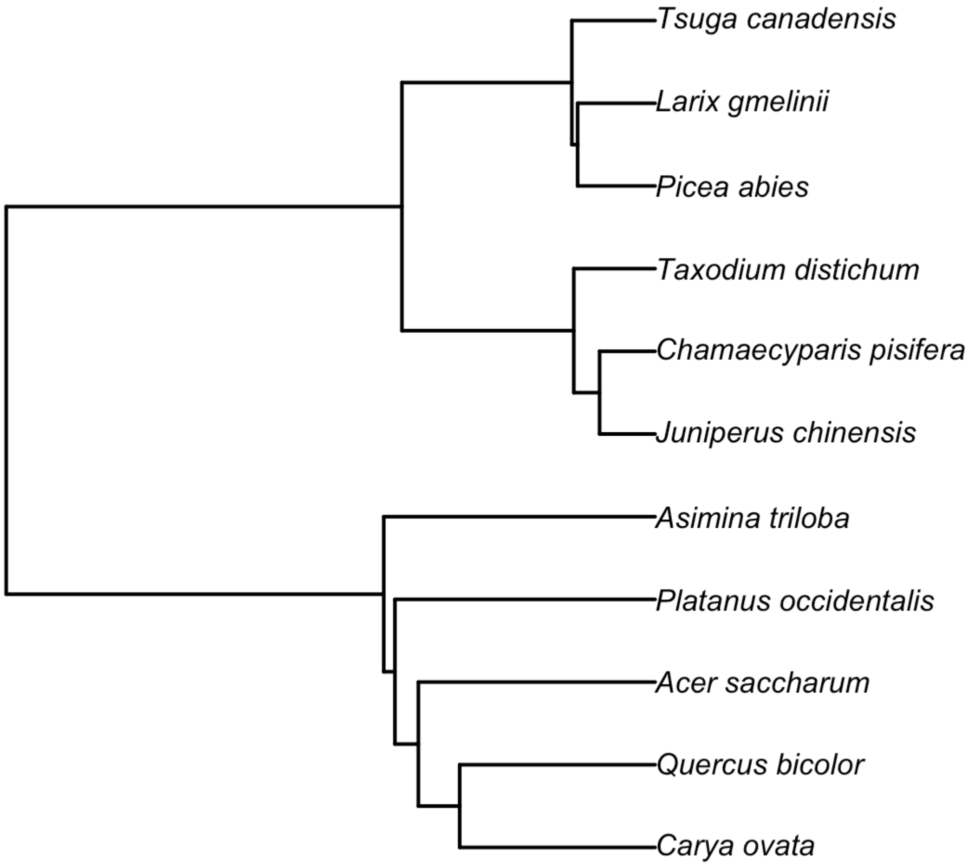


Fig S2) The phylogenetic tree of 11 tree species at The Morton Arboretum, Lisle, IL, USA. The phylogenetic tree was built using R packages: V.PhyloMake, phylo.maker. The generated tree was used for estimation of phylogenetic signals such as Blomberg's K, Lambda, K*, and phylogenetic independent contrasts (PICs). Only Blomberg's K was reported since all signals were consistent throughout the methods.

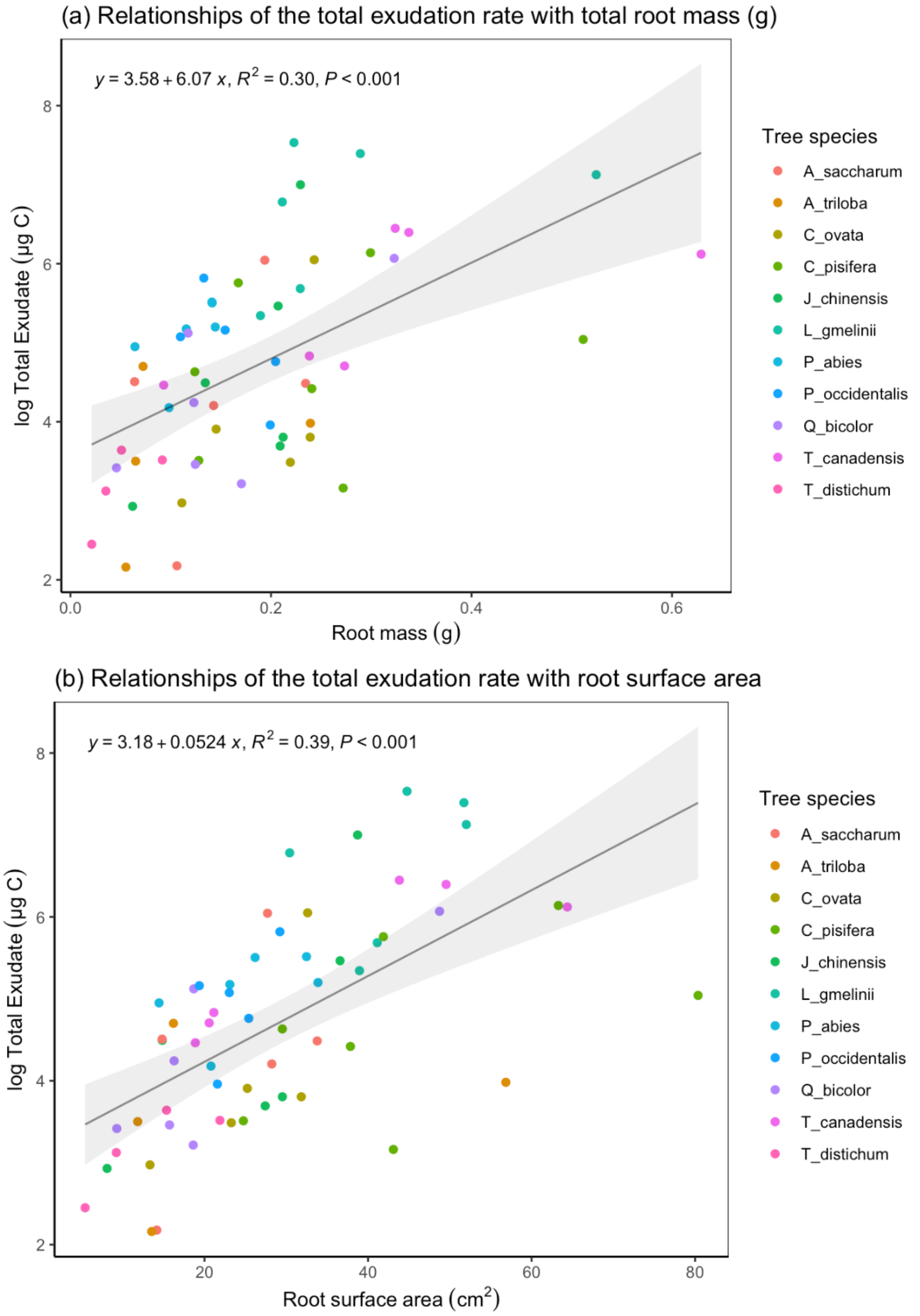


Fig S3) The relationships of the total exudation rate with (a) total root mass (g) and (b) total root surface area ($cm^2$) (bottom) measured for 11 tree species planted at The Morton Arboretum, Lisle, IL, USA. Two figures show the amounts of roots sampled was significantly correlated with total exudate loading (total root mass (g): $R^2 = 0.30$, $p < 0.001$; root surface area ($cm^2$): $R^2 = 0.39$, $p<0.001$). A significant correlation was also found in root length ($R^2 = 0.25$, $p<0.001$), signifying the measurements were robust.

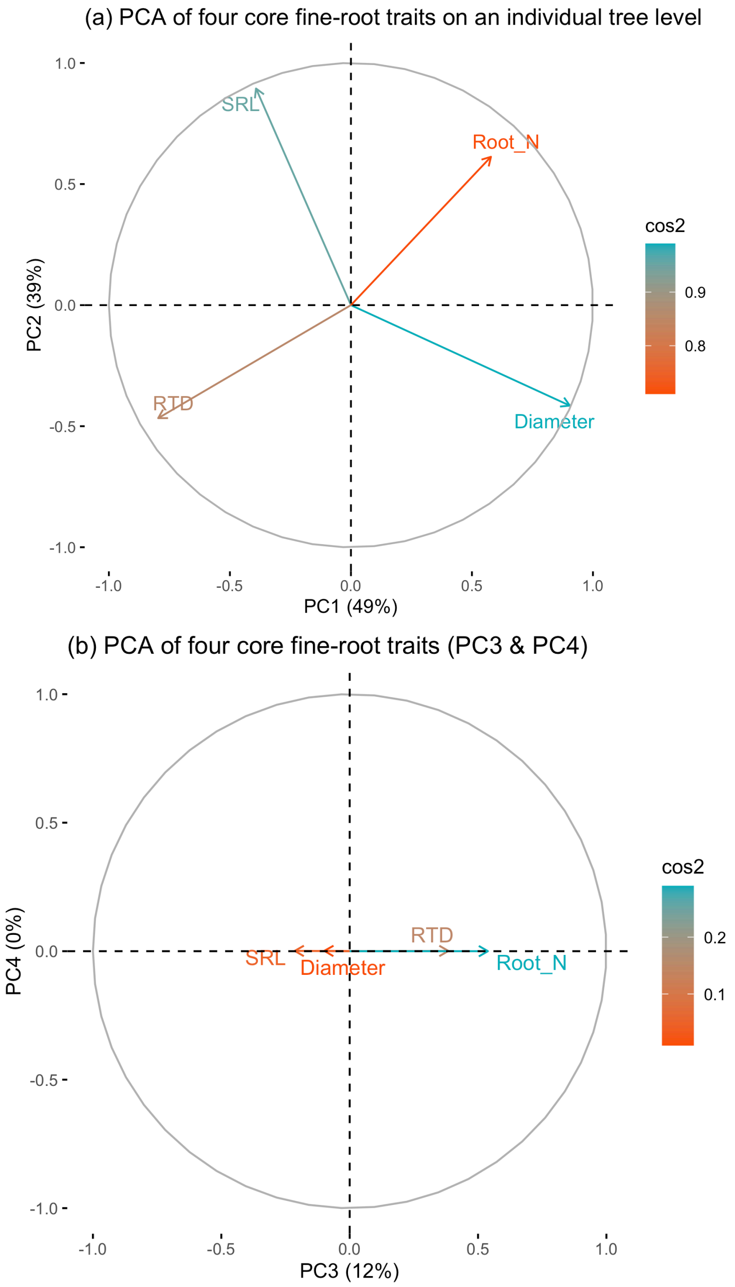


Fig S4) (a) Standardized principal components analysis (PCA) of four core fine-root functional traits, SRL, root diameter, RTD, and root N (Bergmann et al., 2020), measured at The Morton Arboretum, Lisle, IL, USA. Stepwise model selection using redundancy analysis (RDA) identified root tissue density (RTD) as the only significant predictor of exudation rates among root traits (AIC = 3.56, F = 5.05, P = 0.015), as the selected RDA model constrained by RTD explained 8.0% of the total variance in exudation rates, with the remainder attributed to unconstrained variation. Also, note that the total variation explained by two axes were reduced from 84% to 73% from the PCA with seven root traits (Fig 4a). (b) Two additional axes (PC 3 & 4) from the PCA.

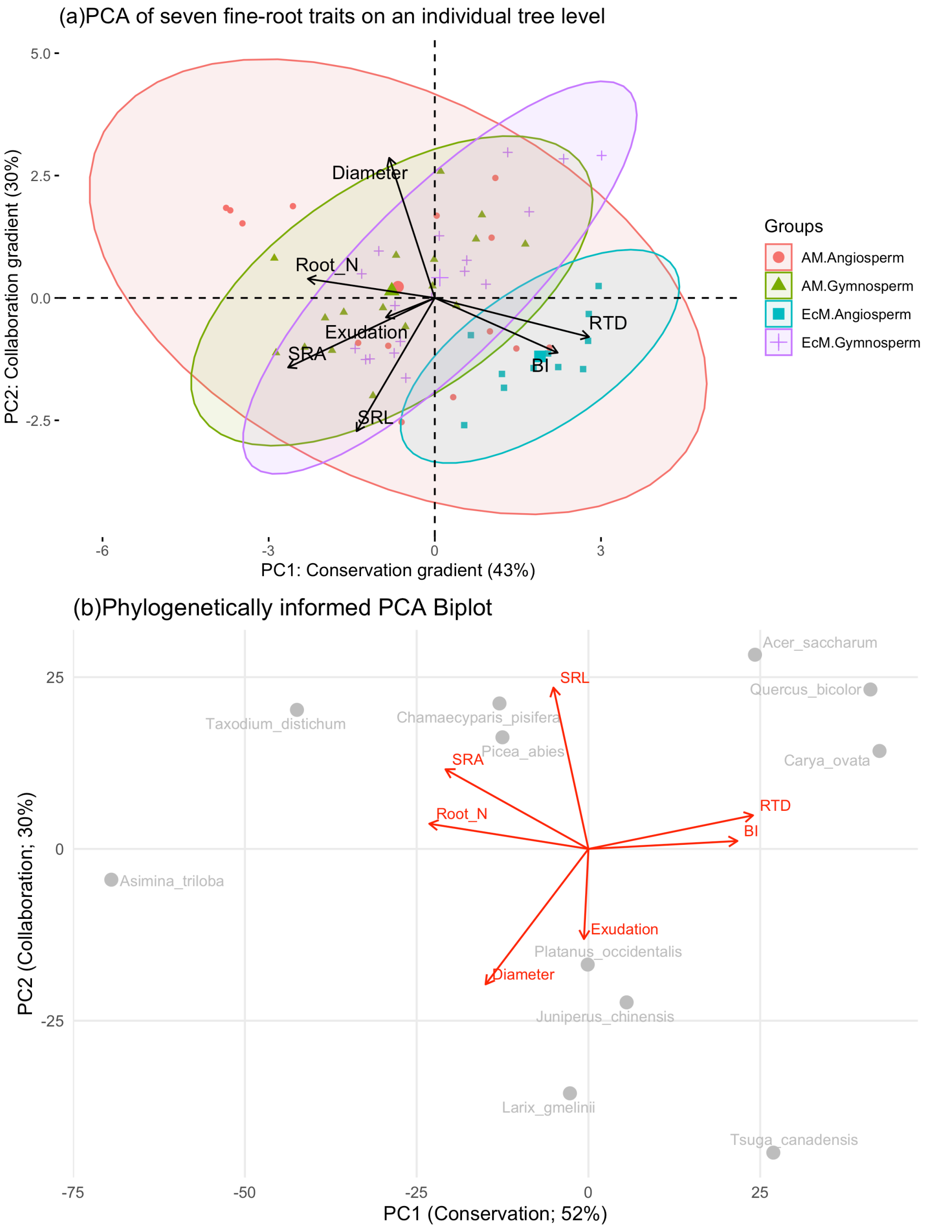


Fig S5) (a) Standardized principal components analysis (PCA) of six core fine-root functional traits, SRL, root diameter, RTD, root N, BI, and SRA measured at The Morton Arboretum, Lisle, IL, USA. (Bergmann et al., 2020). PC1 & PC2 likely represent conservation and collaboration gradient, respectively (Bergmann et al., 2020). Note that the variation explained by two axes were reduced from 84% to 73% (Fig 4a) as exudation rates were added. (b) Phylogenetically informed principal component analysis (pPCA) of root trait variation among 11 mature tree species (n = 11) at The Morton Arboretum, Lisle, IL. The analysis was conducted on species means for seven core root traits using the phytools::phyl.pca function with the "lambda" method to account for phylogenetic structure. While most traits showed weak or nonsignificant phylogenetic signals (Table S2), branching intensity (BI) exhibited a notable phylogenetic pattern (Table S2). Consequently, phylogenetically informed RES axes may capture evolutionary structure but may be less representative than those derived from individual-level data. relatedness and that all traits except BI showed insignificant phylogenetic signals (Table S2).

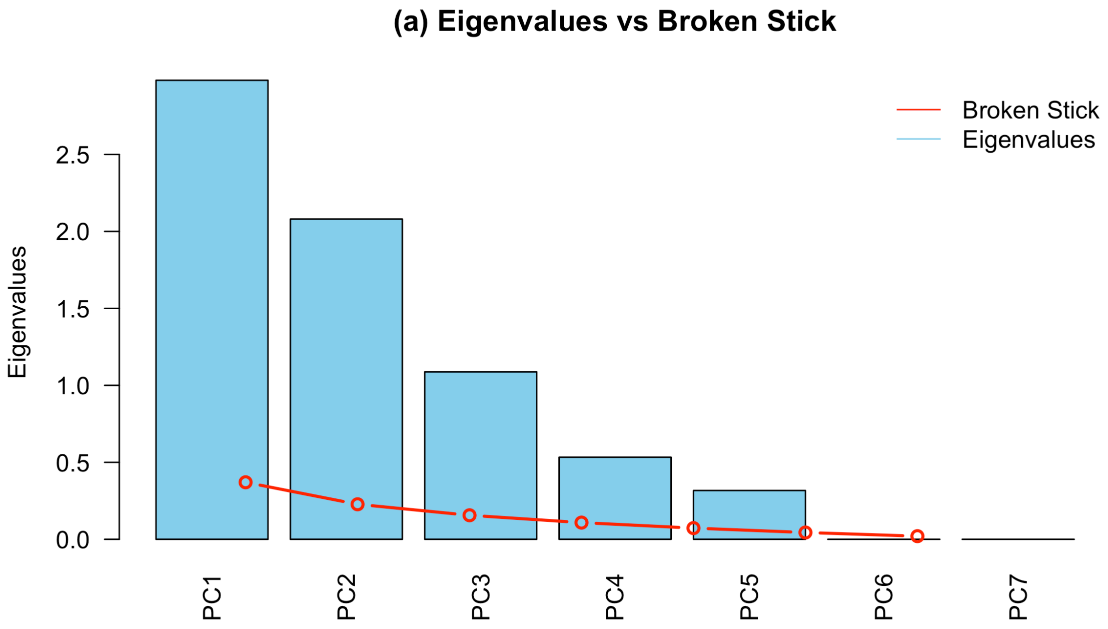


Fig S6) Eigenvalues for the first seven principal components of the root economic space (RES), as projected by standardized PCA of seven root traits including exudation rates (from Fig 4. in the main text). The broken stick model (red circles and lines) is the variance expected under a random distribution of variance, which in all cases is lower than the variance explained by each principal component.

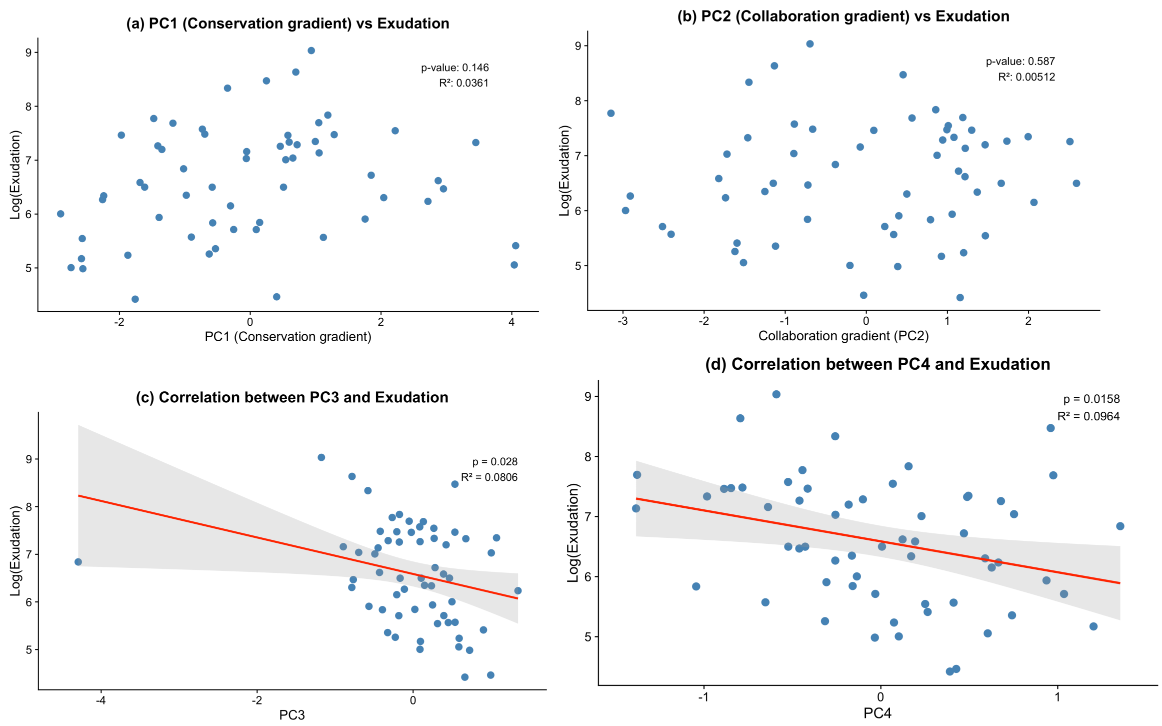


Fig S7) Relationships between root exudation rates and (a) PC1 (conservation gradient), (b) PC2 (collaboration gradient), (c) PC3 (undefined), and (d) PC4 (undefined) of the RES based on six root traits excluding exudation rates (Fig 4). The models show that exudation rates are weakly related to undefined gradients; PC3 ($R^2$ =0.08, p=0.03) and PC4 ($R^2$ =0.10, p=0.02) and unrelated to PC1 and PC2. Notably, considering PC3 and PC4 together (exudation~PC3+PC4) improved the significance of the model ($R^2$ =0.18, p=0.004).

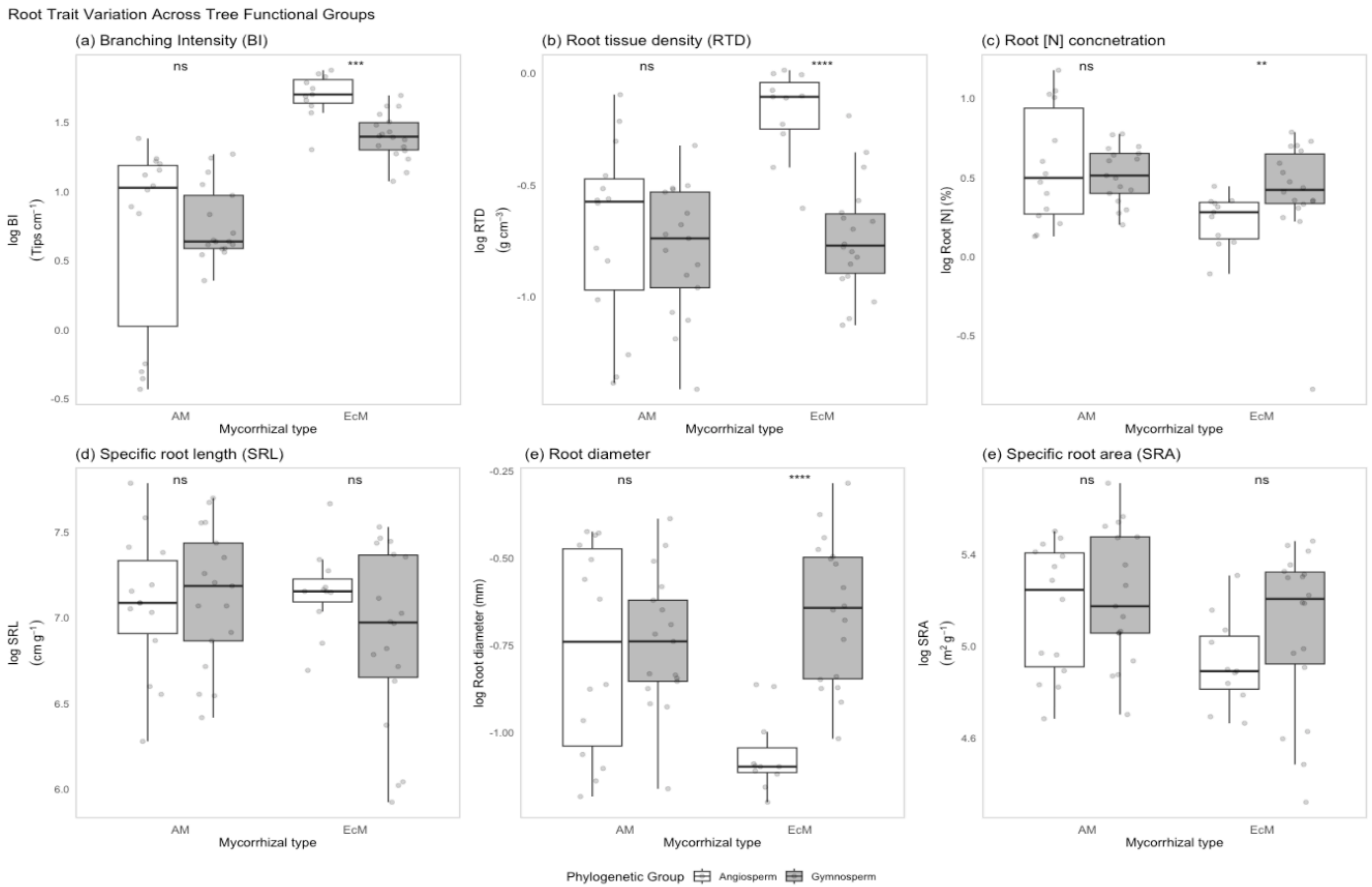


Fig S8) Boxplot of variations of root traits ((a) Branching intensity, (b) RTD, (c) Root N, (d) SRL, (e) diameter, and (f) SRA) measured at The Morton Arboretum, Lisle, IL, USA. Tree species were nested in mycorrhizal types and phylogenetic groups. The central box in each boxplot represents the median and the interquartile range. The whiskers extend to The whiskers extend to the minimum and maximum value. The nested groups were tested with Wilcoxon rank-sum test (non-parametric) is used to compare groups. This test is robust to non-normal data and often used for comparing medians of two groups. Significance levels indicate the significance of difference between angiosperms and gymnosperms nested within mycorrhizal association: ****$p < 0.0001$. *** for $p < 0.001$ ** for $p < 0.01$ * for $p < 0.05$ ·for $p < 0.1$ (marginal significance). Please note that for more robust statistics refer to Table 3 that used the mixed effects models with adjusted p-values from FDR Correction.

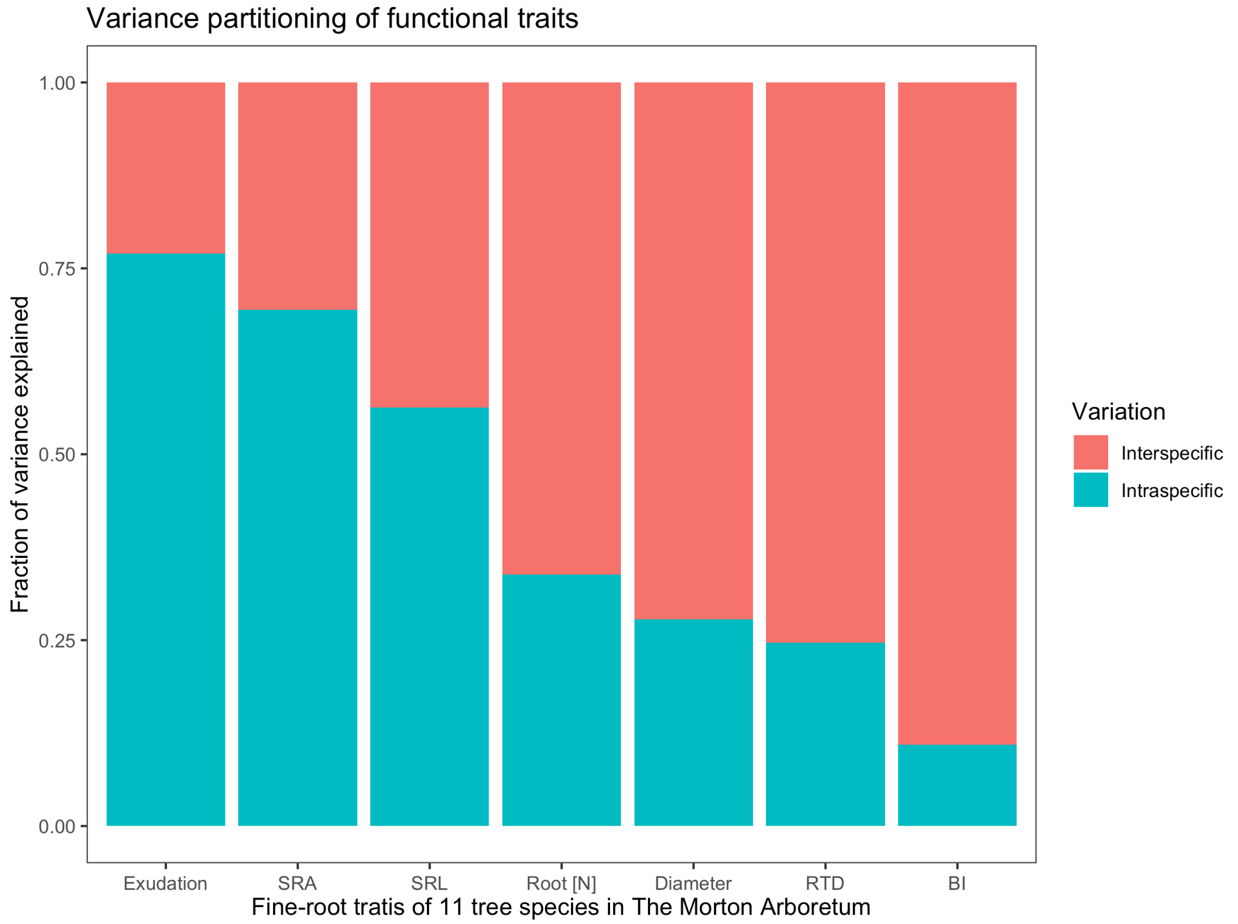


Fig S9) Variance partitioning of fine-root functional traits of 11 tree species. All individuals were mature trees (> 85 years old except *A. triloba*) occurring in monospecific plots at The Morton Arboretum, Lisle, IL, USA. Fraction of variance explained for each root functional trait was calculated from the results of a mixed effect model where monospecific plots were considered as a random effect. N uptake, net inorganic N uptake rate (mg N $d^{-1}$ $g_{root}^{-1}$); Exudation, mass-specific exudation rate (mg C $d^{-1}$ $g_{root}^{-1}$); SRA, specific root area ($cm^2$ $g^{-1}$); SRL, specific root length (m $g^{-1}$); Root [N], root nitrogen concentration (%); Diameter; fine-root diameter (mm); RTD, root tissue density (mg $cm^{-3}$); BI, branching intensity (tips per cm).

**Table S1) Descriptive statistics of the eight fine-root traits of eleven tree species (n=60) in the monodominant forestry plots at The Morton Arboretum, Lisle, IL, USA.**

| Descriptive statistics for root functions & traits | Mass-specific exudation rate (mg C $g_{root}^{-1}$ * $day^{-1}$) | Net N uptake (mg N $g_{root}^{-1}$ * $day^{-1}$) | Net $NH_4^+$ uptake (mg N $g_{root}^{-1}$ * $day^{-1}$) | Net NO3- uptake (mg N $g_{root}^{-1}$ * $day^{-1}$) |
|---|---|---|---|---|
| [a]Min | 0.08 | (973.0) | (450.5) | (690.5) |
| [b]Max | 8.4 | 1,531.9 | 1,113.9 | 702.6 |
| Median | 0.7 | 274.0 | 170.5 | 111.4 |
| Mean | 1.2 | 304.3 | 209.0 | 95.3 |
| SD | 1.4 | 501.3 | 281.8 | 254.6 |
| CV% | 118.8 | 164.7 | 134.8 | 267.1 |

| Descriptive statistics for root functions & traits | Specific root length (m $g^{-1}$) | Root diameter (mm) | Root tissue density (g $cm^{-3}$ ) | N concentration (%) | Specific root area ($cm^2$ $g^{-1}$) | Branching intensity ($cm^{-1}$ ) |
|---|---|---|---|---|---|---|
| [a]Min | 372.5 | 0.3 | 0.2 | 0.4 | 75.3 | 0.7 |
| [b]Max | 2,408.9 | 0.8 | 1.0 | 3.2 | 301.8 | 6.5 |
| Median | 1,212.2 | 0.4 | 0.5 | 1.5 | 171.2 | 3.4 |
| Mean | 1,246.4 | 0.5 | 0.6 | 1.6 | 173.3 | 3.4 |
| SD | 478.1 | 0.1 | 0.2 | 0.5 | 51.0 | 1.5 |
| CV% | 38.4 | 25.4 | 36.5 | 29.9 | 29.4 | 43.9 |

*SD: standard deviation; CV%, coefficient of variation*

[a]Min & [b]Max: the minimum and maximum value of the measured traits across 11 species

**Table S2)** Mixed effects models for the effects of mycorrhizal type on exudation-to-trait relationships. The model considered root traits × mycorrhizal type + (1|species.plot) to predict exudation rates. 'species.plot' represents species-specific monodominant plots at The Morton Arboretum, Lisle, IL. The numbers in the table represent coefficients estimate (Std β) and standard error (Std SE) in brackets. Adjusted p-values were calculated using the Benjamini-Hochberg (BH) correction for multiple testing. Bold value indicates statistical significance of p-values less than 0.05. Significance levels of Adjusted p-values: *p<0.05; **p<0.01; ***p<0.001. The comprehensive table for the effects of phylogeny on trait-exudation relationships is in Table S2.

| **Model to predict exudation rates<br>- AIC values<br>- Marginal & conditional R²** | **Fixed effect term** | **Std β (Std SE)** | **p.value** | **Adjusted p** |
|---|---|---|---|---|
| RTD * Mycorrhizal type | (Intercept) | 6.24 (0.37)*** | **0.000** | **0.000** |
| - AIC=163.2 | RTD | -0.12 (0.45) | 0.784 | 0.896 |
| - $R^2$m = 0.24; $R^2$c= 0.44 | Myco.type [ECM] | -0.53 (0.47) | 0.266 | 0.376 |
| | RTD : Myco.type [ECM] | -2.05 (0.64)** | **0.002** | **0.008** |
| SRA * Mycorrhizal type | (Intercept) | -2.82 (3.32) | 0.400 | 0.533 |
| - AIC=169.3 | SRA | 1.76 (0.64)* | **0.008** | **0.023** |
| - $R^2$m = 0.33; $R^2$c= 0.33 | Myco.type [ECM] | 2.60 (4.49) | 0.565 | 0.714 |
| | SRA : Myco.type [ECM] | -0.35 (0.87) | 0.688 | 0.826 |
| SRL * Mycorrhizal type | (Intercept) | -4.64 (3.20) | 0.152 | 0.243 |
| - AIC=170.7 | SRL | 1.54 (0.45)** | **0.001** | **0.005** |
| - $R^2$m = 0.19; $R^2$c= 0.43 | Myco.type [ECM] | 8.35 (4.45) | 0.065 | 0.131 |
| | SRL : Myco.type [ECM] | -1.10 (0.63) | 0.086 | 0.158 |
| Diameter (mm) * Mycorrhizal type | (Intercept) | 5.22 (0.61)*** | **0.000** | **0.000** |
| - AIC=173.0 | Diameter | -1.49 (0.77) | 0.064 | 0.131 |
| - $R^2$m = 0.10; $R^2$c= 0.27 | Myco.type [ECM] | 2.91 (0.90)* | **0.005** | **0.016** |
| | Diameter : Myco.type [ECM] | 3.09 (1.09)* | **0.010** | **0.027** |
| Root N * Mycorrhizal type | (Intercept) | 6.38 (0.45)*** | **0.000** | **0.000** |
| - AIC=176.6 | Root N | -0.08 (0.72) | 0.917 | 0.945 |
| - $R^2$m = 0.13; $R^2$c= 0.21 | Myco.type [ECM] | -0.04 (0.54) | 0.945 | 0.945 |
| | Root N : Myco.typee [ECM] | 1.34 (0.95) | 0.167 | 0.250 |
| BI * Mycorrhizal type | (Intercept) | 6.36 (0.32)*** | **0.000** | **0.000** |
| - AIC=179.2 | BI | -0.03 (0.36) | 0.941 | 0.945 |
| - $R^2$m = 0.12; $R^2$c= 0.13 | Myco.type [ECM] | 2.88 (1.34) | 0.053 | 0.128 |
| | BI : Myco.type [ECM] | -1.55 (0.93) | 0.120 | 0.206 |

| **Model to predict exudation rates**<br>**- AIC values**<br>**- Marginal & conditional $R^2$** | **Fixed effect term** | **Std β (Std SE)** | **p.value** | **Adjusted p** |
|---|---|---|---|---|
| RTD * Phylogeny | (Intercept) | 5.92 (0.37)*** | **0.000** | **0.000** |
| - AIC=176.5 | RTD | -0.63 (0.55) | 0.264 | 0.511 |
| - $R^2$m = 0.14; $R^2$c= 0.29 | Phylogeny [Gymno] | 0.19 (0.67) | 0.785 | 0.897 |
| | RTD:Phylogeny [Gymno] | -0.30 (0.86) | 0.732 | 0.878 |
| SRA * Phylogeny | (Intercept) | -2.50 (3.64) | 0.494 | 0.698 |
| AIC=171.8 | SRA | 1.72 (0.72) | **0.019** | 0.062 |
| - $R^2$m = 0.21; $R^2$c= 0.43 | Phylogeny [Gymno] | 3.69 (4.63) | 0.429 | 0.664 |
| | SRA:Phylogeny [Gymnosperm] | -0.63 (0.90) | 0.488 | 0.698 |
| SRL * Phylogeny | (Intercept) | -3.12 (3.90) | 0.426 | 0.664 |
| AIC=172.3 | SRL | 1.32 (0.55) | **0.019** | 0.062 |
| - $R^2$m = 0.20; $R^2$c= 0.43 | Phylogeny [Gymnosperm] | 4.93 (4.76) | 0.305 | 0.563 |
| | SRL:Phylogeny [Gymnosperm] | -0.60 (0.67) | 0.374 | 0.641 |
| Diameter * Phylogeny | (Intercept) | 5.66 (0.83)*** | **0.000** | **0.000** |
| AIC=178.7 | Diameter | -0.66 (0.89) | 0.469 | 0.698 |
| - $R^2$m = 0.10; $R^2$c= 0.27 | Phylogeny [Gymnosperm] | 0.68 (1.07) | 0.528 | 0.724 |
| | Diameter:Phylogeny [Gymnosperm] | 0.05 (1.28) | 0.971 | 0.971 |
| Root_N * Phylogeny | (Intercept) | 6.08 (0.40)*** | **0.000** | **0.000** |
| AIC=178.9 | Root_N | 0.36 (0.71) | 0.619 | 0.803 |
| - $R^2$m = 0.10; $R^2$c= 0.24 | Phylogeny [Gymnosperm] | 0.53 (0.54) | 0.339 | 0.602 |
| | Root_N:Phylogeny [Gymnosperm] | 0.11 (0.95) | 0.913 | 0.971 |
| BI * Phylogeny | (Intercept) | 6.26 (0.38)*** | **0.000** | **0.000** |
| AIC=178.0 | BI | -0.02 (0.29) | 0.956 | 0.971 |
| - $R^2$m = 0.13; $R^2$c= 0.16 | Phylogeny [Gymnosperm] | -0.23 (0.65) | 0.722 | 0.878 |
| | BI:Phylogeny [Gymnosperm] | 0.76 (0.53) | 0.174 | 0.364 |

Note) All measured variables were log-transformed to ensure normality of data. (Intercept) represents baseline exudation rate (AM reference). $R^2$m, Marginal $R^2$; $R^2$c, Conditional $R^2$, RTD. $R^2$m represents variance explained by fixed effects only, whereas $R^2$c indicates variance explained by both fixed and random effects. Root Tissue Density (g/cm3); SRA, Specific Root Area (m2/g); Specific Root Length (cm/g); Diameter, root mean diameter (mm); Root N, Root N Concentration (%); BI, Branching Intensity (cm-1).

**Table S3)** Variance partitioning of fine-root functional traits

| | Percent variance explained | |
|---|---|---|
| **Root functions and traits** | **Within-species (-plot)** | **Across-species (-plot)** |
| Mass-specific exudation rate | 77% | 23% |
| Specific root length (SRL) | 56% | 44% |
| Root diameter | 28% | 72% |
| Root tissue density (RTD) | 25% | 75% |
| Root [N] concentration | 34% | 66% |
| Specific root area (SRA) | 69% | 31% |
| Branching intensity (BI) | 11% | 89% |

Note) Within-species (-plot): intraspecific percent variance explained, Across-species (-plot): interspecific percent variance explained

**Table S4)** Best predicting Mixed effects models of fine-root morphological traits on mass-specific exudation rate

| | *Dependent variable:* | | |
|---|---|---|---|
| | Mass-specific exudation rate | | |
| | Large model | Intermediate model | Selected model |
| Mycorrhizal type | -0.10 | | -0.09 |
| | (0.39) | | (0.38) |
| Phylogenetic type | -0.18 | | -0.18 |
| | (0.33) | | (0.33) |
| SRA | 1.14$^{**}$ | 1.45$^{**}$ | 1.13$^{**}$ |
| | (0.42) | (0.45) | (0.39) |
| Root [N] | -0.05 | 0.36 | |
| | (0.43) | (0.50) | |
| Interaction term | 1.33$^{**}$ | | 1.32$^{**}$ |
| | (0.48) | | (0.48) |
| Constant | -6.35$^{**}$ | -7.95$^{***}$ | -6.34$^{**}$ |
| | (2.10) | (2.29) | (2.03) |
| Observations | 60 | 60 | 60 |
| Log Likelihood | -75.40 | -80.56 | -75.41 |
| Akaike Inf. Crit. | 166.81 | 171.13 | 164.82 |
| Bayesian Inf. Crit. | 183.56 | 181.60 | 179.48 |
| Significance levels | $^{*}p<0.05$; $^{**}p<0.01$; $^{***}p<0.001$ | | |

All measured variables were log-transformed to ensure normality of data. For the units in each variable, refer to Table 2. The numbers in the table represent coefficients estimate and standard error in brackets.

Note) The best predicting root traits for exudation rates were selected based on AIC and BIC. Variables were selected from the predictors chosen by a multiple linear regression, and cross-validated by the mixed effects model, where the procedure considered each monodominant species-plot as a random effect, considering that each plot comprised the same tree species with a similar age group (90-100 years), and the days of data collection campaigns were randomly chosen and blocked by the plot. The larger model considered myco + phylo + SRA + root[N] + myco:phylo, the intermediate model considered SRA and root[N], and the selected model contained SRA + myco:phylo.

**Table S5)** Results of principal component analysis (PCA) on the six root traits, mass-specific exudation rates from 11 tree species measured at The Morton Arboretum, Lisle, IL, USA. The table includes the proportion of variation explained (top section) and loading scores of traits on each component (bottom section)

| Component | Eigenvalue | Difference | Proportion | Cumulative |
|---|---|---|---|---|
| 1 | 2.98 | 0.90 | 42.60 | 0.43 |
| 2 | 2.08 | 0.99 | 29.72 | 0.72 |
| 3 | 1.09 | 0.55 | 15.54 | 0.88 |
| 4 | 0.53 | 0.22 | 7.62 | 0.95 |
| 5 | 0.32 | 0.32 | 4.53 | 1.00 |
| 6 | 0.00 | 0.00 | 0.00 | 1.00 |
| 7 | 0.0 | | 0.00 | - |

| Vairable | PC 1 | PC2 | PC3 | PC4 |
|---|---|---|---|---|
| Exudation | -0.17 | -0.09 | **0.87** | -0.27 |
| Diameter | -0.15 | **0.64** | 0.18 | 0.18 |
| SRL | -0.26 | **-0.61** | -0.09 | 0.09 |
| RTD | **0.52** | -0.18 | -0.15 | -0.36 |
| Root[N] | **-0.43** | 0.09 | -0.20 | -0.82 |
| BI | **0.42** | -0.25 | 0.38 | -0.04 |
| SRA | **-0.50** | -0.32 | 0.03 | 0.28 |

Note) PCs with eigenvalue close to or greater than 1 are considered as significant (Tabachnik & Fidell, 1996). Variable loading scores greater than 0.4 were considered significant, and significant traits to contribute each PC appear in bold. Exudation, mass-specific exudation rate (µgC d-1 g.root-1); SRA, specific root area (cm2 g−1); specific root length (m g−1); Root [N], root nitrogen concentration (%); Diameter; fine-root diameter (mm); RTD, root tissue density (mg cm−3); BI, branching intensity (tips per cm).
